# Integrated single-cell whole genome sequencing and spatial transcriptomics reveal latent intra-tumoral heterogeneity in ovarian cancer

**DOI:** 10.1101/2025.10.08.676897

**Authors:** Rania Bassiouni, Yuxin Jin, Lee D. Gibbs, Jing Qian, Solomon O. Rotimi, Heather Miller, Michelle G. Webb, Seeta Rajpara, Javier Arias-Stella, David W. Craig, Lynda Roman, John D. Carpten

## Abstract

The mortality rate of ovarian cancer remains disproportionately high compared to its incidence. This is partly due to a high level of intra-tumoral heterogeneity that promotes disease recurrence and treatment failure. In this study, we describe degrees of heterogeneity revealed by single-cell whole genome sequencing and spatial transcriptomics of five epithelial ovarian carcinomas. At the cellular level, we describe pseudo-diploid cells that match the malignant cell population in both somatic variant and copy number patterns. At the clonal and subclonal levels, we describe diversification associated with copy number gains and whole genome doubling. In multi-clonal samples, we infer evolutionary relationships from single cell copy number, loss of heterozygosity analysis, and somatic variant detection, and correlate these with tissue histology and gene expression programs. In one sample, we identify functionally consequential copy number alterations that contribute to molecular diversity, cell proliferation, and inflammation in a minor clone that persisted without major expansion alongside a more complex major clone. In another, we describe a complex evolutionary history including a spontaneous reversion of a driver mutation in a secondary clone, which correlated with a switch in oncogenic expression programs.

## Introduction

Ovarian cancer is the deadliest gynecological malignancy in the United States, with more than half of patients diagnosed with late-stage disease [1]. Most ovarian cancers are epithelial in origin and can be classified into four major histological subtypes: serous, endometrioid, clear cell, and mucinous [2]. Among these, high grade serous ovarian cancer (HGSOC) is the predominant histological subtype. It is also the most aggressive and has the poorest overall prognosis, with the majority of patients experiencing disease relapse after an initially favorable response to chemotherapy [3]. It is estimated that up to 90% of HGSOC patients diagnosed with metastatic disease will develop total treatment resistance [4].

In contrast to HGSOC, clear cell ovarian cancer (CCOC) is less common and often presents in younger patients at early stage, during which it has a more favorable prognosis than HGSOC [5, 6]. However, CCOC patients with late-stage disease have poorer responses to platinum-based chemotherapy and poorer progression-free survival rates than HGSOC patients [6, 7]. Thus, both HGSOC and CCOC are focuses of efforts to improve ovarian cancer outcomes.

In addition to unique histology and clinical characteristics, HGSOC and CCOC are biologically distinct, as evidenced by several studies that have characterized the genomic landscapes of HGSOC and CCOC [8–13]. Despite nearly ubiquitous *TP53* mutations, HGSOC exhibits fewer somatic mutations than cancers originating in other tissues [14, 15]. Instead, HGSOC is characterized by high-level aneuploidy and recurrent copy number alterations (CNA), which are often not directly therapeutically actionable [15]. CCOC exhibits a greater diversity in somatic driver variants, dominated by mutations in *ARID1A* and *PIK3CA*, but with *TERT* promoter, *KRAS*, *TP53*, *ATM*, *CTNNB1*, and *PTEN* mutations also occurring at appreciable frequencies [12, 16]. Recurrent CNAs are also present in CCOC, although with a lower degree of disorder than that seen in HGSOC [13].

Epithelial ovarian cancers, and particularly HGSOC, are recognized for their intra-tumoral heterogeneity, which is reflected in both CNA and mutational profiles obtained from multi-region and longitudinal tumor sampling [17–19]. Spatial transcriptomics (ST) has been utilized to infer clonal compositions in HGSOC and associated biology in histological context [20]. However, copy number inferred from gene expression data cannot be assessed with confidence at high resolution. Single-cell DNA sequencing (scDNA-seq) provides the most direct method to assess CNA heterogeneity and resolve subtle clonal structures that would be otherwise obscured [21–24], but scDNA-seq alone does not describe tumor biology.

In this study, we integrate scDNA-seq with ST analysis of five ovarian cancer samples to assess biological manifestations of clonal diversity. From scDNA-seq, we assess copy number and somatic mutations at the single cell and cluster level to identify clonal CNAs, whole genome doubling (WGD) events, and pseudo-diploid cells. In multi-clonal samples, we infer tumor evolution in the context of CNAs, driver mutations, and loss of heterozygosity. From matched ST data, we demonstrate that gene expression programs controlling metabolism, cell proliferation, and inflammation correspond to clone-specific CNAs identified in scDNA-seq. We also report, for the first time, a spontaneous reversion of an oncogenic *CTNNB1* driver mutation in a secondary clone and associated compensatory gene expression programs. These cases highlight the insights that can be gained from integration of high-resolution genotypic and phenotypic information.

## Methods

### Samples

Tissue specimens were obtained as OCT-embedded frozen blocks from the University of Southern California Keck School of Medicine Gynecological Tissue and Fluid Repositories under a protocol approved by the Institutional Review Board (#HS-18-00948). The repository also provided matched peripheral blood buffy coat where available. Patients provided written informed consent for sample collection. Samples were all epithelial ovarian cancers (EOC) with varied histological subtypes: three HGSOC, one CCOC, and one predominantly CCOC mixed with endometrioid. Subtype information was obtained from pathology reports and confirmed by contemporary review. All tumors sampled were high stage (≥ IIIC) and high grade. All samples were assessed for tumor adequacy and contained >60% tumor by area in regions sampled. Sample information is summarized in Supplemental File 1.

### DNA extraction and whole exome sequencing

Genomic DNA was extracted from buffy coat using the DNeasy Blood & Tissue Kit (Qiagen) according to the manufacturer’s protocol. Tumor DNA was extracted from ten 10-micron scrolls of frozen tumor tissue using the AllPrep Mini Kit (Qiagen) according to the manufacturer’s protocol. DNA quantity and quality was assessed to ensure sufficiency for library preparation. Following enzymatic fragmentation of the DNA, dual indexed sequencing libraries were prepared using the SureSelect Low Input Target Enrichment System with SureSelect XT Human All Exon V6 target enrichment probes (both Agilent Technologies). Pooled libraries were sequenced at 2×150 on a NovaSeq 6000 instrument (Illumina). Raw sequencing data was demultiplexed using bcltofastq (v.2.20.0.422), and FASTQs were aligned to human reference genome GRCh38 using bwa mem (v 0.7.17) to generate binary alignment map (BAM) files.

### Bulk copy number analysis

The sequenza R package (v 3.0.0) was used to process the paired tumor/normal BAMs against a reference genome track (GRCh38) and generate allele-specific copy numbers [25]. A wiggle track file for the reference genome was first created using a window of 50 bp. Sequenza then processed the tumor/normal pair by tabulating reads per genomic bin, followed by segmentation to identify regions of the genome with aberrant copy number values. Sequenza output was used to plot copy number, depth ratios, and allele frequencies across the genome.

### Nuclear isolation for single cell DNA sequencing

Single nuclei were isolated from frozen tissue sections as recommended by 10x Genomics (Document CG000167, Rev A). Briefly, tissue sections were thawed on ice, lysed, and physically homogenized by pestle. Following a dual step centrifugation, the recovered nuclei were washed in PBS with 0.04% BSA and passed through a 40-micron strainer to remove aggregates. Nuclei were stained with ethidium homodimer-1 and counted with a Countess II FL cell counter (Invitrogen).

### Single cell DNA sequencing (scDNA-seq)

Single nuclei were processed with Chromium Single Cell DNA Reagent Kits (10x Genomics) according to the manufacturer’s protocol (Document CG000153, Rev B-C). Briefly, individual nuclei are partitioned into single droplets of a hydrogel matrix, within which the nuclei are lysed. This droplet containing denatured genomic DNA is then co-encapsulated with a gel bead containing hexamer primers and a unique 16 nucleotide barcode, allowing amplification and unique indexing of DNA from individual cells. The encapsulation is then dissolved, and barcoded DNA is pooled and used to generate Illumina-compatible sequencing libraries. Following quality assessment, libraries were sequenced at 2×100 on a NovaSeq 6000 instrument (Illumina).

### Single cell copy number (CN) calling and metrics

Sequencing data was processed with the Cell Ranger DNA pipelines (v. 1.0.0, 10x Genomics). Demultiplexing was performed with *cellranger-dna mkfastq* and was followed by *cellranger-dna cnv* for alignment to human reference genome GRCh38, cell identification, and copy number estimation. The number of cells detected in the samples ranged from 532 to 1437. An average of ∼1.6B mapped, de-duplicated cell-associated reads were processed per sample, for a mean of ∼1.75M reads per cell, sufficient for detecting copy number event sizes of ∼1 MB. The median effective reads per MB ranged from 420 to 786. scDNA-seq metrics per sample can be found in Supplemental File 1.

### Postprocessing of single cell CN calls

To generate a high quality dataset, we filter on both CN calls and on cells in a multi-step fashion. First, noisy cells and low quality calls were removed from the data set (*Filtering of single cell CN calls*, below). The remaining cells were subject to clustering analysis to reveal major copy number profiles (*Clustering and subclustering of CN profiles*). Clusters comprised of degraded DNA, likely representing apoptotic cells, were identified and removed. Tumor copy number profiles were then used to identify and remove possible doublets and computational artifacts (*Removal of doublets and artifacts*). Remaining cells were then subject to a final round of clustering and subclustering.

### Filtering of single cell CN calls

Cells were classified as noisy and removed from the data if they met either of 2 criteria: (1) ploidy confidence of the cell was low, or (2) the DIMAPD score of the cell – a measure of bin-to-bin variation – was higher than the threshold equivalent to a p-value of 0.1 when a Gaussian distribution was fit to the data. This DIMAPD filtering is more stringent than the default performed by cellranger, to provide confidence that we were removing cells in the process of active DNA replication, which may confound further analysis. In the remaining cells, individual CN calls per 20KB bin were assessed for quality and calls with quality scores < 15 were removed. Finally, regions of the genome with mappability < 90% were removed. Following this process, the clean data was organized in a matrix of CN calls per 20 KB bin, per cell barcode.

### Clustering and subclustering of CN profiles

We employed a clustering strategy based on the one described by Velazquez-Villarreal et al [26]. CN calls for 500 sequential bins were aggregated to generate a mean ploidy value at 10 MB resolution in each cell, using the GenomicRanges R package (v. 1.50.2) [27]. The cells were then subject to maximum likelihood genetic clustering using the R package adegenet (v. 2.1.10) [28]. Data was clustered at all values of *k* up to *k*=20, and the Bayesian Information Criterion (BIC) and Akaike Information Criterion (AIC) were evaluated. Clustering solutions selected based on BIC reflected underclustering, while AIC better captured heterogeneous populations (Supp. Fig. 1). AIC was therefore used to select *k* for clustering. Cells in each cluster were then subject to a second round of clustering by the same method to determine subclusters. Barcodes retained in each sample and their cluster assignments can be found in Supplemental File 3. Discriminant analysis of principal components (DAPC) was performed using the adegenet package to assess the relationships between determined clusters and subclusters. Euclidean distance of cell pairs within clusters was calculated with the amap R package (v. 0.8-19).

### Removal of doublets and artifacts

Following a first round of clustering, tumor clusters which represented a pre-WGD profile were identified and used to generate baseline copy number profiles. OV511 and OV594 contained two baseline profiles representing distinct clones. A profile representing a diploid-tumor doublet was then generated by adding 2 across the baseline tumor profiles. In OV511 and OV594, a tumor-tumor doublet profile was also generated by summing both baseline profiles. Cells matching the doublet profiles were removed from the dataset. We then removed cells that may represent artifacts of copy number overfitting by ensuring that cells contained a 10 Mb region with an odd copy number, following the strategy of McPherson et al [24]. Final assignments and metrics for each barcode per sample can be found in Supplemental File 2.

### Determination of haplotype-specific single cell CN

From scDNA-seq, the diploid cell cluster was identified in each sample and used as a matched-normal for germline variant calling as follows. The sample BAM file was used to create a mini-BAM containing only diploid cells by adapting the script provided by Velazquez-Villareal et al [26]. The diploid mini-BAM was used to generate a germline variant call format (VCF) file using GATK HaplotypeCaller (v4.1.8.0) [29]. Eagle2 (v2.4.1) [30] was then used to estimate haplotype phase, and autosome heterozygous single nucleotide polymorphisms (SNPs) were identified using bcftools (v1.17) [31]. The Copy-number Haplotype Inference in Single-cell by Evolutionary Links (CHISEL) algorithm [32] was then used to compute phased single-cell copy number. Single-cell read depth ratio (RDR) was calculated across 10Mb bins, and B-allele frequency (BAF) was calculated across 50Kb bins for each sample. Cells previously classified as noisy (see ‘Filtering of single cell CN calls’) were removed manually from the RDR and BAF datasets, followed by CHISEL-calling of allele- and haplotype-specific copy numbers in remaining cells. Loss of heterozygosity (LOH) was inferred from phased haplotypes. We consider as LOH any region that lacks representation of one allele, which may manifest as hemizygosity, copy-neutral LOH, or LOH with high-level gain of the remaining allele. For assessment of BAF at the cluster level, average BAF at identified SNP positions was binned into 100Kb blocks to smooth out the noisiness inherent to single cell data.

### Somatic variant calling

Somatic single nucleotide variants, insertions, and deletions were identified from scDNA-seq at the cluster level using Strelka2 (v2.9.2) in tumor-normal mode [33]. Using mini-bams (see ‘Determination of haplotype-specific CN’), tumor clusters were assessed against matched diploid clusters. Somatic variant calling was performed with default parameters against the GRCh38 reference genome. Variants were annotated using bcftools (v1.14) against the dbSNP build 154 database. Variant effects were annotated and converted to Mutation Annotation Format using vcf2maf (v1.6.22) with the VEP database [34, 35]. Variants predicted to be pathogenic were visually inspected using IGV (v2.14.1) [36].

### CN visualization

Single cell CN was visualized on heatmaps, where each row corresponded to a cellular barcode, and each column corresponding to a 10 MB bin using the ComplexHeatmap R package (v. 2.14.0) [37]. The presence or absence of LOH, as determined by CHISEL, was similarly visualized. Heatmaps were organized by and annotated with cluster and subcluster information. Psuedobulk copy number line plots were created with R package ggplot2 (v. 3.4.2) [38] and reflect the average ploidy value at each 10 MB bin, with shaded regions depicting standard deviation where appropriate. Copy number line plots paired with chromosome ideograms are displayed at 1 Mb resolution and were created with R package karyoploteR (v. 1.24.0) [39].

### Phylogenetic analysis

Phylogenetic analysis was performed based on the strategy employed by Minussi at al [21]. Pairwise Manhattan distance was calculated on copy number values per cell (in 1Mb bins) or per subcluster (mean value at 20Kb bins), using the amap R package. Phylogenetic analysis was then performed with the balanced minimum evolution (ME) algorithm in the R package ape (v. 5.7-1) [40]. Trees were rooted by specifying a diploid cell or (sub)cluster as the outgroup, then visualized with R package ggtree (v. 3.6.2) [41]. For visualization purposes, the diploid clusters were displayed in negative distance space.

### Visium spatial transcriptomics

Archival, mounted and H&E stained FFPE tissue was utilized with the Visium Cytassist Spatial Gene Expression assay (10x Genomics). Hardset coverslips were first removed as described in 10x Genomics Demonstrated Protocol CG000518 (Rev C). The Visium assay was then performed following manufacturer’s protocol, using 11mm x 11mm capture slides. Following quality assessment, libraries were sequenced (2×100) on an Illumina NextSeq2000 instrument, to generate a minimum of 25,000 read pairs per spot under tissue. Full metrics can be found in Supplemental File 1.

Raw sequencing data was processed with the *spaceranger mkfastq* and *spaceranger count* pipelines (v. 3.0.1, 10x Genomics) as described previously [42]. Human reference genome GRCh38 was used for alignment. Raw count data was imported into the R package Seurat (v. 4.3.0.1) and was normalized with the SCTransform method from the R package sctransform (v. 0.3.5) [43, 44].

### CN inference from spatial transcriptomics data

The R package infercnv (v. 1.18.1) [45] was used to infer copy number profiles as follows. Normalized count data was first subject to ESTIMATE analysis using the R package estimate (v. 1.0.13) [46] to determine a tumor purity score for each spot. By comparing tumor purity scores with histological annotation, we determined that a tumor purity score of 0.7 marked a reasonable threshold to distinguish tumor spots. Therefore, spots with a tumor purity score < 0.7 were considered reference spots, while spots with tumor purity > 0.7 were considered tumor, and provided to inferCNV as observations. Raw counts were then analyzed with inferCNV in subcluster mode, with HMM-determined CN predictions, to identify copy number events in the observation spots relative to the reference spots. Tumor subgroups were determined by inferCNV using the Leiden algorithm.

### Gene set enrichment analysis (GSEA)

GSEA and single sample GSEA (ssGSEA) were performed using GenePattern software [47] as previously described [42]. Analysis was performed against the Hallmark gene sets [48] and custom positional gene sets. Cell cycle phase determination was predicted with Seurat’s gene set-based CellCycleScoring feature.

### Data and code availability

All datasets generated in this study will be made available in an appropriate repository. Code necessary to reproduce the analysis will be made available in the supplemental files.

## Results

### Characteristics of clinical samples

We obtained frozen tumor tissue from 5 EOC cases. Samples OV440, OV594, and OV773 are HGSOC; sample OV150 is of mixed CCOC and endometrioid histology; and sample OV511 is CCOC. We selected samples with similar stage and grade reflective of clinically challenging late-stage disease. We sampled cases with diverse patient self-reported race: three patients were Hispanic/Latino (OV440, OV511, and OV773), one was Asian (OV150), and one was non-Hispanic White (OV594). All patients were younger than 63, the median age at diagnosis for ovarian cancer [1]. While we did not select based on *BRCA* status, we had prior knowledge of a germline *BRCA2* E49X mutation in sample OV440. All samples were treatment-naïve and obtained from primary biopsies or surgeries.

### Benchmarking of copy number calling across modalities

To evaluate subclonal copy number heterogeneity, we subjected isolated nuclei from each sample to individual DNA barcoding and shallow whole-genome sequencing (single-cell DNA sequencing, scDNA-seq) as described in Methods. Approximately 1.7 million deduplicated reads were sequenced per cell, for an average of 541 reads per megabase (Supp. File 1). Copy number data and metrics produced from the Cell Ranger DNA pipeline were used to filter out low quality data. Filtered data was then subjected to maximum likelihood genetic clustering analysis to resolve clusters and subclusters.

In sample OV440, we resolve four clusters (Fig. 1A). Cluster 1 is comprised mainly of diploid cells, cluster 2 is the baseline tumor, and clusters 3 and 4 likely represent subclonal whole genome doubling events, which occur nearly ubiquitously in HGSOC [24]. Subclusters within cluster 2 were distinguished by minor CN differences in chromosomes 1, 2, 8 13, and X (Fig 1A). Utilizing the CHISEL algorithm [32], we determined corresponding single-cell haplotype specific copy numbers, revealing extensive loss of heterozygosity (LOH), including the copy-neutral LOH of the entirety of chromosome 17, encompassing *TP53*, and at chromosome 13q12, encompassing *BRCA2* (Fig 1B).

**Figure 1.**
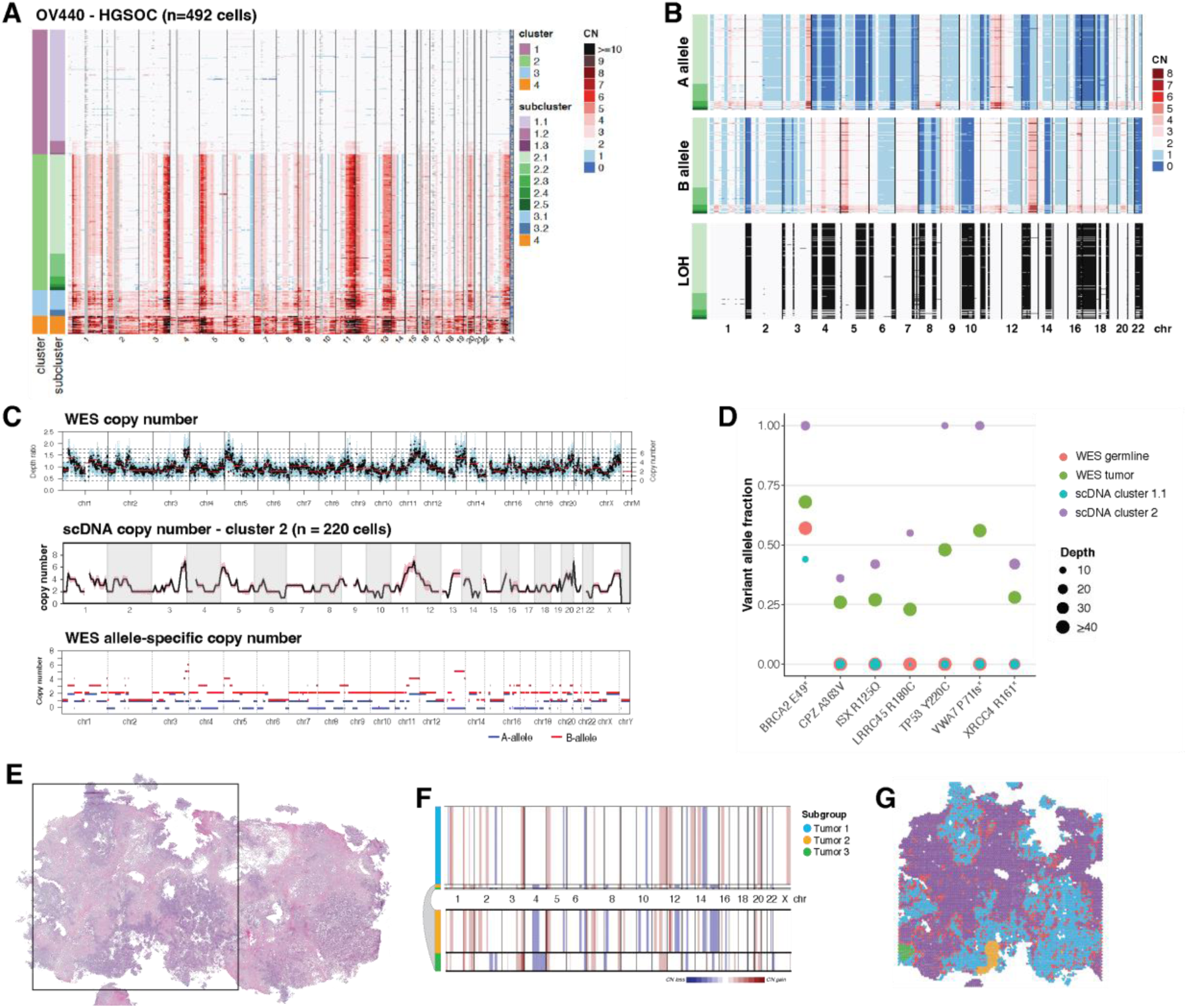
Benchmarking of copy number determination across modalities. (A) Copy number (CN) as determined by single cell whole genome sequencing of sample OV440, binned at 10 Mb resolution. Each row displays data from a single cell. Heatmap colors represent absolute copy number, while gray regions represent unmappable areas, or low quality calls that were removed from the analysis. Chromosomes are delineated and annotated on the x-axis. Cells are ordered by cluster and subcluster assignment on the y-axis, which were determined by successive rounds of maximum likelihood genetic clustering. (B) Single cell allele-specific copy numbers for OV440 cluster 2, as determined by CHISEL. Loss of heterozygosity (LOH) is inferred from the A- and B-allele profiles as described in Methods. LOH is represented by black shading. (C) Copy number as determined by whole exome sequencing (WES) of OV440 is shown in the top panel. The middle panel displays the average copy number of cluster 2 from scDNA-seq, binned at 10 Mb resolution. The shaded region around the line represents the standard deviation. The bottom panel is the allele specific copy number determined from WES. (D) The variant allele fractions of select germline and somatic mutations in OV440 as determined from both WES and scDNA-seq. The size of the bubbles represents the number of reads spanning the mutation locus in each sample. (E) H&E stained tissue of sample OV440. The square represents the 11mm x 11mm region sampled by spatial transciptomics. (F) CNAs inferred from ST using inferCNV. Subclusters were determined by Leiden clustering and are annotated on the left bar. HMM-based CN prediction was then performed at the subcluster level.Three malignant subgroups were identified; the smaller groups are expanded below the plot for clarity. (G) Subgroups from (F) mapped spatially. Spots labeled Interface were determined to be mixtures of malignant and diploid cells (Supp. Fig 2).

We benchmarked our scDNA data against copy number called from bulk whole exome sequencing (WES). WES-derived total copy number and allele-specific copy number matched the profile of cluster 2, confirming that WGD cells are a minor population in this tumor (Fig 1C). We also identified somatic variants from scDNA-seq using diploid subcluster 1.1 as a surrogate matched normal to tumor cluster 2. Variants included the expected *TP53* mutation, as well truncating and nonsynonymous mutations in several other genes (Fig. 1D). Variants determined from scDNA-seq were verified in WES, with variant allele fractions (VAF) reflecting that WES sampled a mixture of malignant and non-malignant cells (Fig. 1D). The germline *BRCA2* truncating mutation and somatic *TP53* Y220C mutation both have a VAF of 1 determined from scDNA-seq, consistent with LOH at both these loci.

We next assessed the fidelity of copy number inferred from ST of OV440 to the ground truth of scDNA-seq (Fig. 1E-G). Application of the inferCNV algorithm revealed three subgroups with distinct copy number profiles. The largest of these matched the profile of scDNA-seq cluster 2, although inferCNV could not differentiate between total copy numbers of 2 and 3, or detect WGD. The two other subgroups corresponded to spatially constrained, histologically unique, and transcriptionally distinct groups of cells that likely represent true subclones that were not sampled in scDNA-seq (Supp. Fig 2). Thus, copy number inferred from ST can generally be matched to scDNA-seq, minding the constraints of sampling bias and algorithm sensitivity.

### Intratumoral copy number heterogeneity in EOC

We similarly assessed single cell copy number profiles of the four additional EOC samples (Fig. 2). All samples contained one diploid cluster, at least one tumor cluster, and at least one WGD cluster. Tumor clusters were often subdivided into multiple subclusters, approximating tumor clones and subclones.

**Figure 2.**
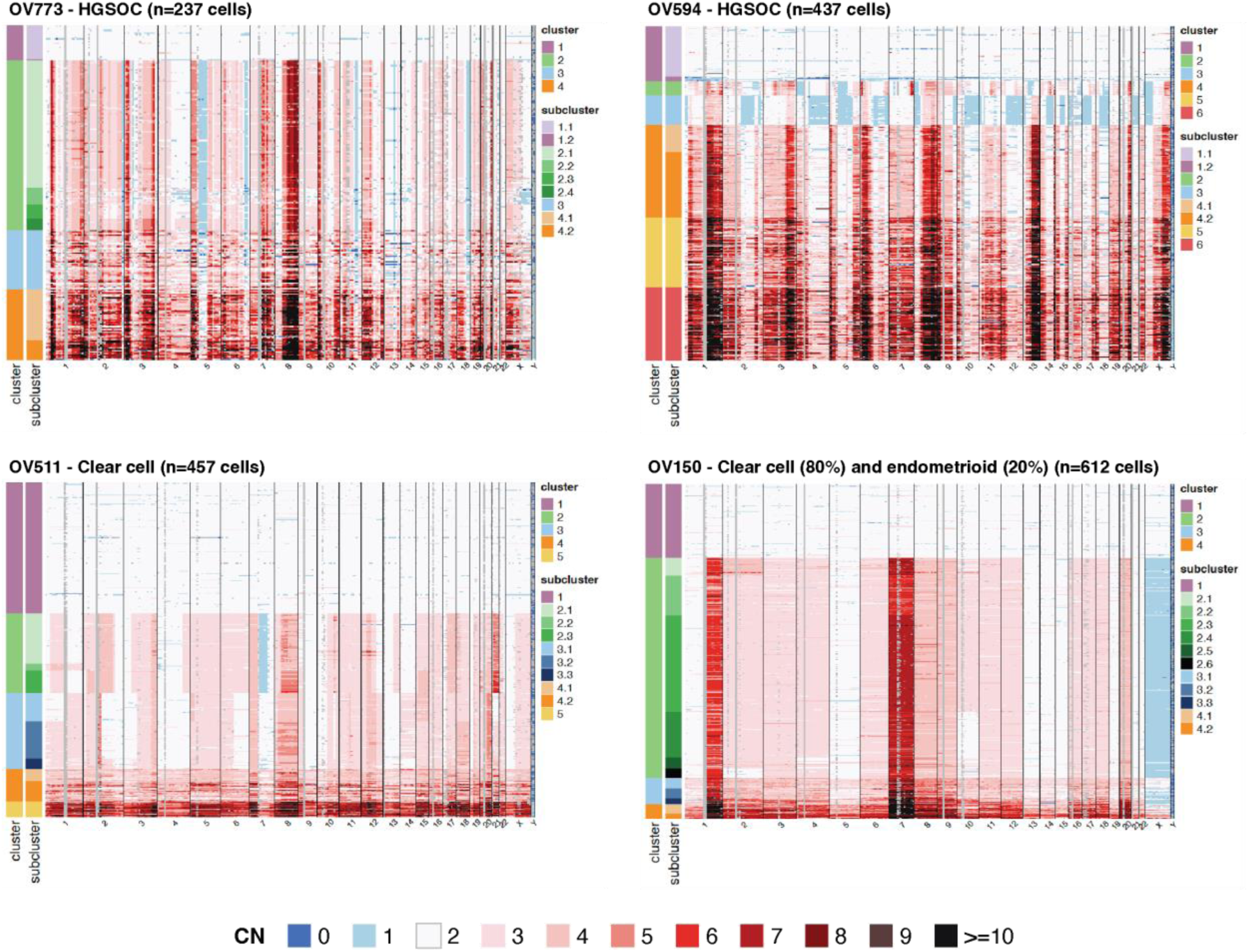
Intratumoral copy number variation in additional EOC samples. Copy number (CN) as determined by single cell DNA sequencing of HGSOC samples OV773 and OV594, and CCOC samples OV511 and OV150. Copy number is displayed at 10Mb resolution. Cluster and subcluster assignments are noted on the left axis.

The HGSOC samples OV773 and OV594 both exhibited highly disordered genomes and harbored somatic clonal *TP53* mutations with chr17 LOH (Supp. Fig. 4). OV511 was characterized by arm-level and segmental CNAs, while OV150 was characterized by aneuploidy and a lack of focal events, likely reflecting mitotic disjunction. OV150 also harbored a clonal *KRAS* G12V mutation (Supp. Fig. 4). OV773 and OV150 each contained one clone (cluster 2), and OV594 and OV511 each contained two distinct clones with further subclonal diversity (clusters 2 and 3). We describe the latter two samples in further detail subsequently in this report.

We assessed the relationships between identified clusters using discriminant analysis of principal components (DAPC) (Fig. 3A). Baseline ploidy was a primary contributor to intra-sample heterogeneity, with WGD driving a main discriminant function. To examine diversity within clusters, we calculated all unique pair-wise distances between individual cells within each cluster assignment (Fig. 3B). Once again, and consistent with recent findings [24], WGD clusters exhibited the greatest intra-cluster diversity. Interestingly, the greatest intra-cluster variability occurred in regions of the genome with copy number gains, regardless of baseline ploidy (Fig. 3C). These results confirm that ovarian cancers exhibit significant intra-tumoral and intra-clonal heterogeneity, primarily driven by copy number gains and WGD events.

**Figure 3.**
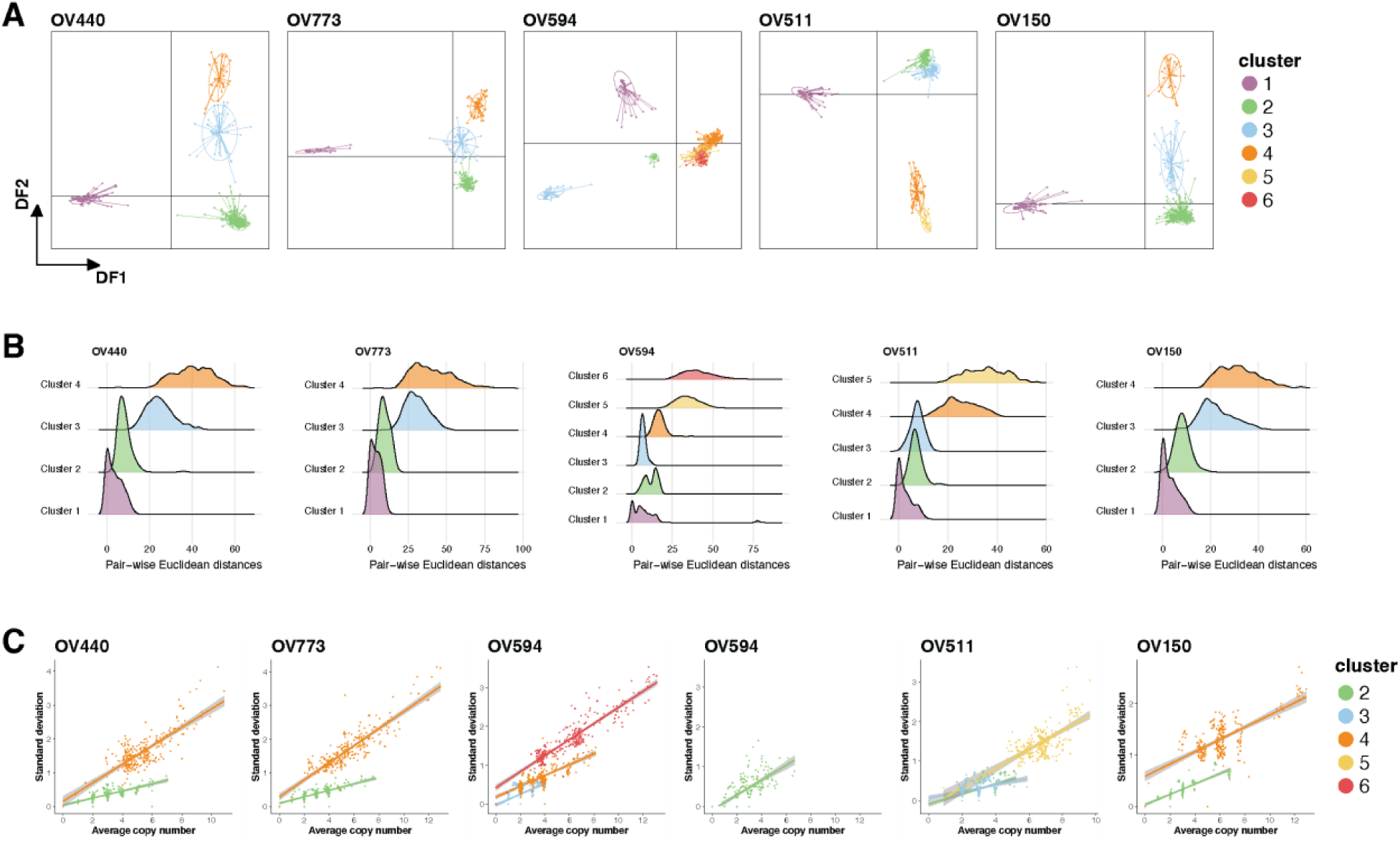
Assessment of cluster diversity. (A) Discriminant analysis of principal components (DAPC) was utilized to describe relationships between clusters in each sample. Plots illustrate discriminant functions (DF) 1 and 2, and points represent individual cells. (B) Ridgeplot displaying all possible pair-wise (cell-cell) Euclidean distances within each cluster and sample. (C) The relationship between mean copy number and standard deviation per 10Mb genomic bin in tumor clusters. OV594 cluster 2 is shown separately for clarity. Linear regression with shaded confidence interval are displayed.

### Assessment of pseudo-diploid cells

In all samples profiled, diploid clusters contained individual cells with sparse and low-level CNA. In OV440, these cells were abundant enough to comprise their own subcluster, 1.2 (Fig. 1A). Such “pseudo-diploid” cells have been previously described with varied interpretations. Early scDNA-seq experiments uncovered pseudo-diploid cells that did not match the CN profiles of the corresponding malignant cells and therefore seemed unrelated [49, 50]. However, a later study described pseudo-diploid cells with non-random and progressive CN patterns that converged to that of the malignant profile [51]. In two of our samples, we find evidence for the latter scenario.

In OV440, subcluster 1.2 harbors CN gains at non-random sites that match regions that are further amplified in tumor cluster 2 (Fig. 4A). DAPC and phylogenetic analysis confirm that subcluster 1.2 is distinct from the true diploid subcluster 1.1, but is still a great distance from cluster 2 (Fig. 4B-C). Subcluster 1.2 also does not appear to harbor any LOH, including at the *TP53* locus (Supp Fig. 5), suggesting that it represents a very early form of malignant transformation. Due to low sequencing depth, we were unable to determine whether these cells harbored the clonal *TP53* Y220C mutation. However, one cell from subcluster 1.2 was found to harbor a truncating frameshift mutation in *VWA7* (Fig 4D), which we had identified as a somatic variant of high VAF in this sample (Fig. 1D).

**Figure 4.**
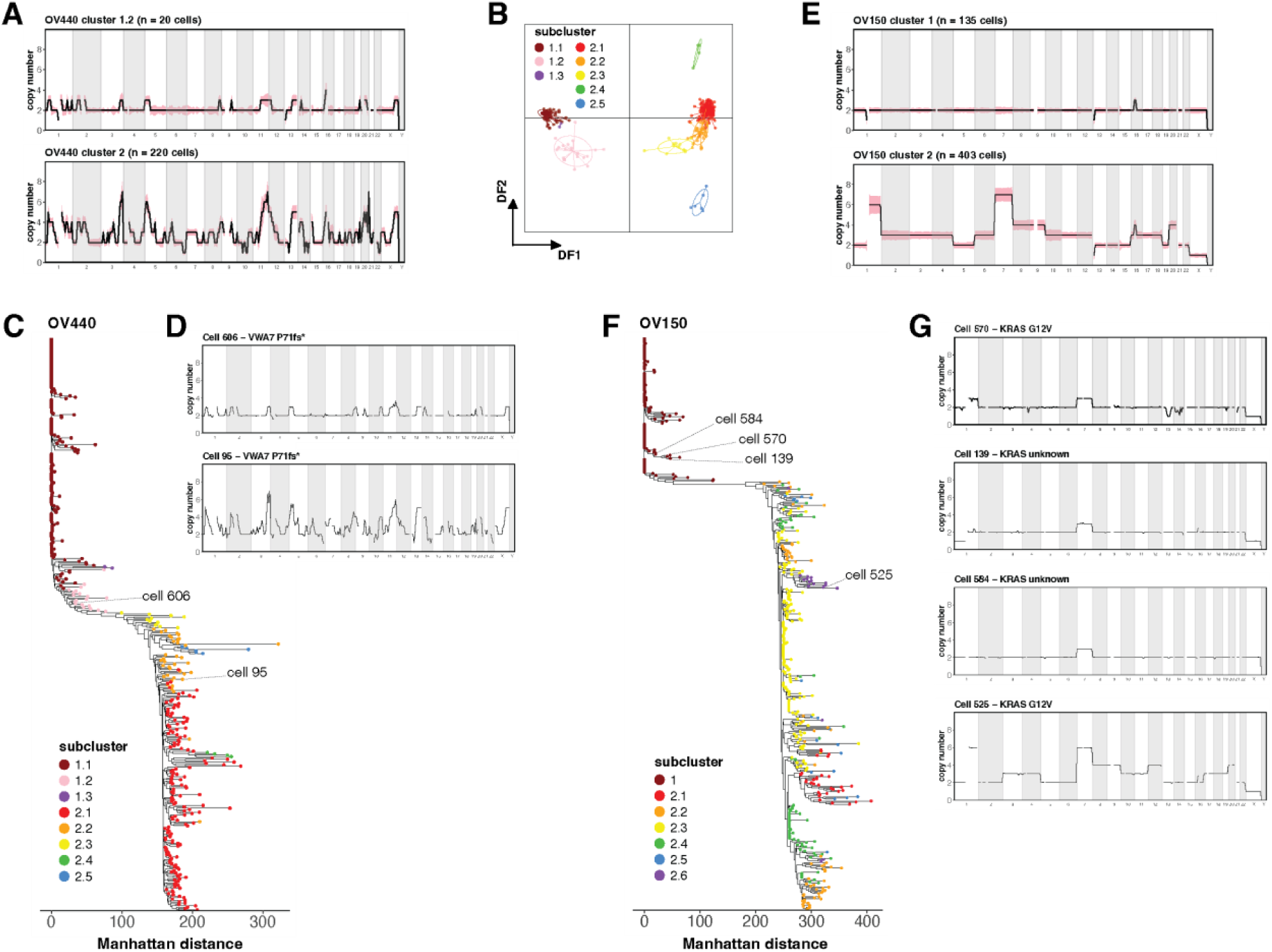
Assessment of pseudo-diploid cells in OV440 and OV150. (A) Average copy number at 10 Mb resolution across the genome for subcluster 1.2 and cluster 2 in sample OV440. Shaded region represents standard deviation. (B) DAPC analysis applied to the subclusters of OV440. (C) Single cell phylogeny of OV440, based on Manhattan distance and rooted at a true diploid cell. Tips represent cells colored by subcluster assignment. (D) The copy number profile of two single cells annotated in (C) that harbor a *VWA7* frameshift truncating mutation. Cell 606 belongs to pseudo-diploid cluster 1.2. Cell 95 is a malignant cell representative of cluster 2. (E) The average copy number profiles of clusters 1 and 2 from OV150. (F) Single cell phylogeny of OV150. (G) Copy number profiles of single cells annotated in (F). Cell 570 is a pseudo-diploid cell with a *KRAS* G12V mutation, while cells 139 and 584 are pseudo-diploid cells for which KRAS status could not be ascertained. Cell 525 is a representative malignant cell with a *KRAS* G12V mutation.

**Figure 5.**
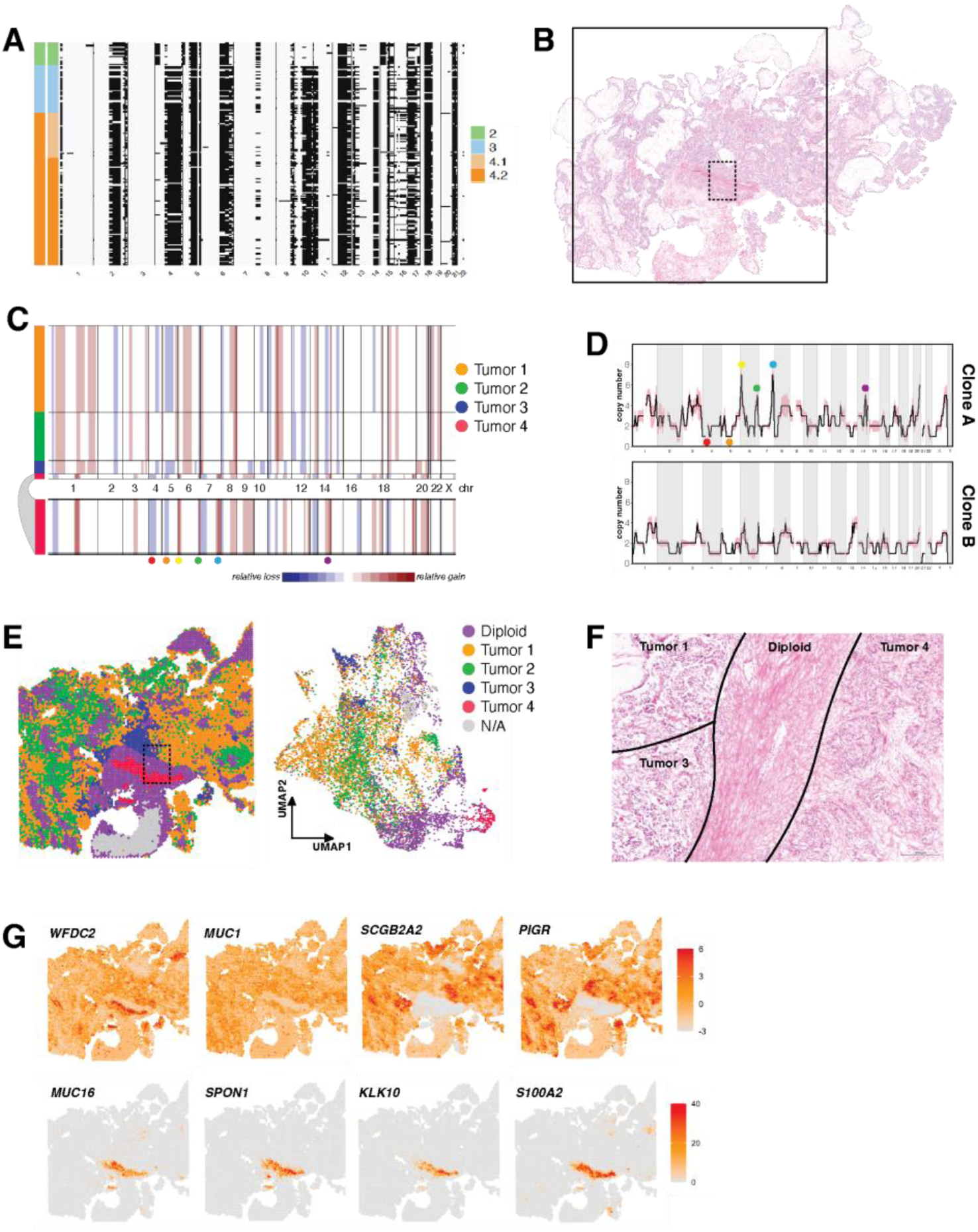
Distinct transcriptional profiles in multi-clonal HGSOC. (A) Single cell LOH inferred from allele-specific copy number for tumor clusters of OV594. LOH is represented by black shading. (B) H&E stained tissue section of sample OV594. The solid square denotes the 11mm x 11mm area assessed by ST. (C) Copy number inferred from ST data by inferCNV. Tumor 4 is expanded beneath the plot for clarity. (D) Average copy number of OV594 clusters 2 and 3 derived from scDNA-seq. The shaded area around the line indicates the standard deviation. The colored landmarks correspond to those indicated in (C) and help match Tumor 4 to Clone A. (E) Spatial map and UMAP plot of the subclusters determined by inferCNV and shown in (C). Diploid spots were those used as reference and showed no evidence of CNA and are comprised mostly of stromal cells. Spots labeled N/A did not contain sufficient transcript density and were filtered out by inferCNV. (F) A magnified and rotated region corresponding to the dotted rectangle in (B) and (E) labeled with inferCNV assignments. Scale bar corresponds to 200 microns. (G) Spatially mapped expression for HGSOC-related genes. Scale bars represent Pearson residuals of expression.

In OV150, no pseudo-diploid subcluster was resolved, but several cells within cluster 1 had apparent CNA, particularly in chr7 (Fig. 2). Notably, chr7 also had the greatest magnitude of gain in the malignant cells (Fig. 4E). Phylogenetically, these pseudo-diploid cells demonstrated grouped divergence from diploid cells (Fig. 4F). Resolving the *KRAS* G12V variant revealed one pseudo-diploid cell in cluster 1 that harbored the mutation (Supp. Fig 3, Fig 4G). This cell had a gain of chr 1q and 7, and loss of X. Similar CNA patterns were seen in other pseudo-diploid cells for which *KRAS* status could not be ascertained (Fig 4G). Thus, in both OV440 and OV150, pseudo-diploid cells appear related to malignant cells in both somatic variants and CNA site preference.

We also note an interesting observance in the single-cell phylogeny of OV150. While most subclusters emerge from single or closely grouped branching events, subcluster 2.4 appears to emerge independently at several different positions (Fig. 4F). Subcluster 2.4 is characterized by two copies of chr10, with most other malignant cells having 3 copies (Fig 2). Closer examination reveals considerable variation at chr1 and chr7 within subcluster 2.4 (Supp Fig. 6). Thus, subcluster 2.3 may not represent a true discrete subclone in the evolutionary sense, but illustrates the dynamic nature of CNA accumulation in cells prone to aneuploidy.

**Figure 6.**
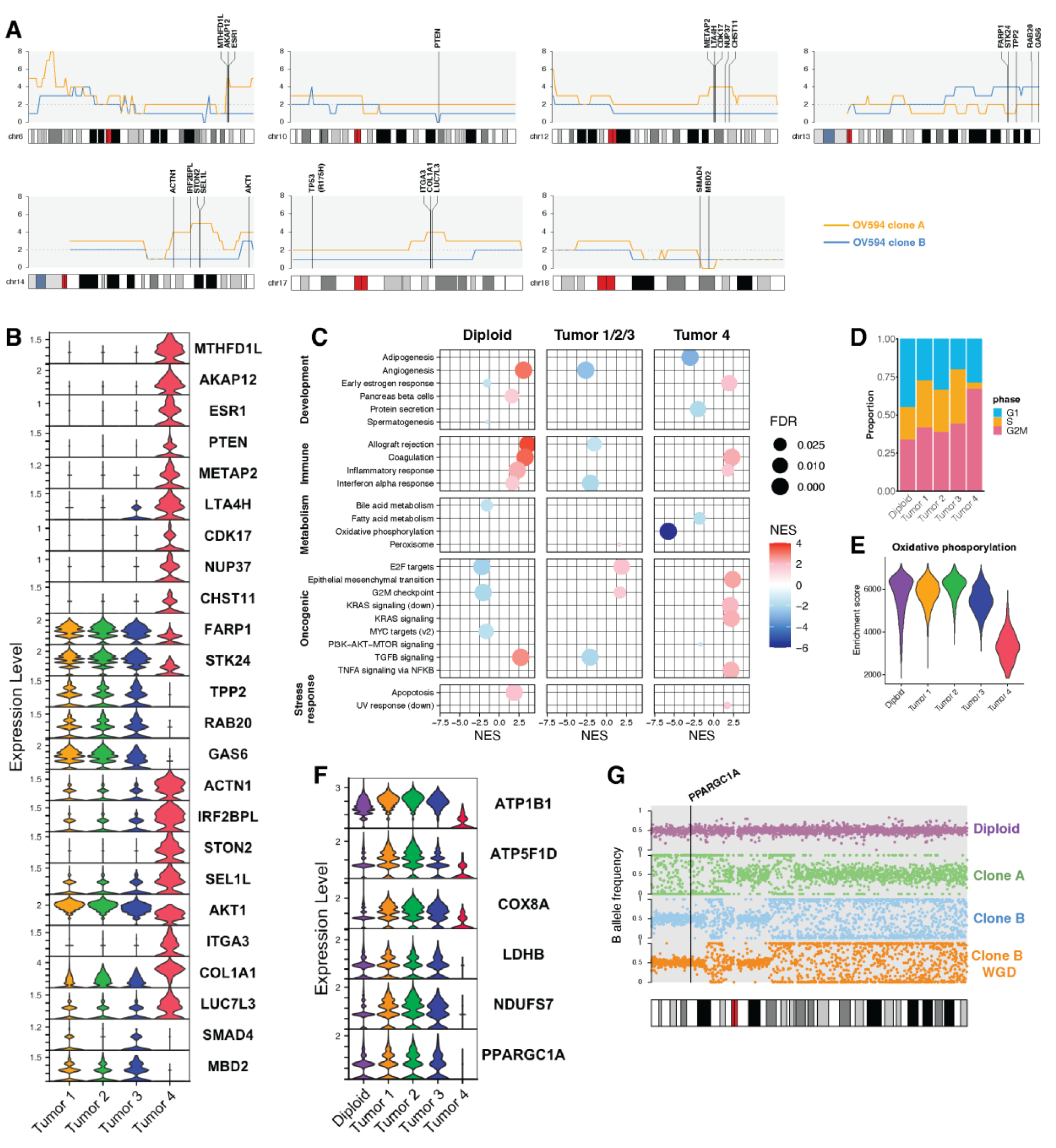
Consequential CNA in a HGSOC minor clone. (A) Averaged copy number of OV594 clones A and B derived from scDNA-seq for chromosomes 6, 10, 12, 13, 14, 17, and 18. Plots are shown at 1Mb resolution, and selected genes are annotated at their genomic positions. (B) Log-transformed expression of genes annotated in (A) in the 4 CN-defined tumor subgroups determined in Fig 6. Tumors 1, 2, and 3 correspond to Clone B, and Tumor 4 corresponds to Clone A. (C) Gene set enrichment analysis was performed against the 50 Hallmark gene sets. Only results with a normalized enrichment score (NES) > 1.5 and a FDR q-value < 0.05 are displayed here. (D) Cell cycle states of spots assigned to tumor subgroups as a proportion of the total number of spots in each group. (E) Summary of per-spot enrichment scores in each tumor subgroup for the Hallmark oxidative phosphorylation gene set, as determined by single sample GSEA. (F) Log-transformed gene expression of representative nuclear-encoded oxidative phosporylation genes. *PPARGC1A* encodes the transcriptional coactivator PGC-1α. (G) B allele frequency in 100 Kb blocks across chromosome 4 for diploid cells, Clone A, and Clone B at baseline and after WGD. These correspond to scDNA-seq clusters 1, 2, 3, and 4, respectively. A BAF of 0.5 reflects allelic balance and heterozygosity, while BAFs of 0 and 1 reflect LOH. The genomic location of *PPARGC1A* is annotated.

### Distinct transcriptional profiles in a multi-clonal HGSOC

Two distinct malignant copy number profiles were resolved in OV594 (Fig. 2); we refer to these as Clone A and Clone B moving forward. We consider Clone A (cluster 2, n = 19 cells) the minor clone, while Clone B (cluster 3, n =38 cells) is likely the dominant clone and undergoes two rounds of WGD (cluster 4, n = 121 and cluster 6, n = 96). Importantly, we found no evidence for WGD of clone A. DAPC suggests that clone A is wholly distinct from clone B (Fig. 3A). However, both clones harbor a *TP53* R175H mutation (Supp. Fig 3) and several regions of shared LOH (Fig. 5A). We therefore infer that they arose from the same ancestor, but with divergent evolutionary trajectories.

To understand the biological relevance of the distinct clones, we assessed histology and ST data for OV594 (Fig 5B). InferCNV predicted four distinct CN profiles from ST data (Fig. 5C). We refer to these inferred copy number profiles as Tumors 1 through 4, to distinguish them from scDNA-seq clusters. Tumors 1, 2, and 3 shared similar CN profiles and corresponded to the profile of Clone B, while Tumor 4 was unique and matched the profile of Clone A (Fig 5C-D).

When plotted spatially and in dimensionally-reduced space, Tumor 4 occupied a discrete location adjacent to, but not intermixed with, the other tumor groups (Fig 5E). Tumor 4 also exhibited a solid growth pattern, while the region corresponding to Tumors 1, 2, and 3 had a distinct papillary growth pattern (Fig. 5F). We examined the expression of known ovarian cancer biomarkers in the tumor groups (Fig. 5G). All tumor regions expressed *WFDC2* and *MUC1*.

However, only Tumors 1, 2, and 3 expressed *SCGB2A2* and *PIGR*. Interestingly, only Tumor 4 expressed *MUC16*, also known as CA125, the most widely used clinical diagnostic serum marker for ovarian cancer [52]. Tumor 4 also exclusively expressed *SPON1*, *KLK10*, and *S100A2* – all have been proposed as prognostic markers in EOC [53–55].

### Identifying consequential CNA in a HGSOC minor clone

We next examined whether unique CN alterations in the minor clone of OV594 may be driving functional consequences. From scDNA-seq, we identified several regions of CNA in Clone A that were associated with gene expression differences in ST data. Most of these regions were associated with CN gains, with two notable exceptions: Clone B was found to harbor a focal deep deletion of *PTEN* with clone-specific LOH, while Clone A harbored a deep deletion of 18q21.2, containing *SMAD4* and *MBD2* (Fig 6A). Loss of gene expression was confirmed in the corresponding ST subgroups (Fig. 6B).

Clone B harbored a gain of a gene-dense portion of chr13, including genes linked to cell cycle regulation (*TPP2*, *RAB20* [56, 57]), cell survival (*GAS6*, *STK24* [58, 59]), and cell motility (*FARP1* [60]). Copy number gain was reflected in increased ST gene expression (Fig. 6B). Among Clone A-specific gains is an amplification of 6q25.1, containing *ESR1* – an uncommon event in ovarian cancer that is associated with strong protein positivity and potential response to anti-hormone therapy [61]. Gene set enrichment analysis (GSEA) also revealed a supportive significant enrichment of early estrogen response genes in tumor 4 (Fig. 6C).

Other regions of gain in Clone A contained genes linked to cell proliferation and invasion (*METAP2*, *ACTN1*, *ITGA3*, *COL1A1, NUP37* [62–66]), drug resistance (*AKAP12*, *COL1A1 [67, 68]*), and inflammation (*ITGA3*, *LTA4H [69, 70]*). These genes and related gene sets were significantly enriched in Tumor 4, including programs for inflammatory response and epithelial mesenchymal transition (EMT) (Fig. 6C). Tumor 4 was also comprised of a greater proportion of cells in G2/M phase (Fig. 6D). Cell cycle arrest at either G1 or G2/M has been linked to EMT in cancer and other pathologies [71–73].

GSEA analysis also revealed an absence of oxidative phosphorylation in Tumor 4 (Fig. 6C, E). Assessment of gene expression of individual electron transport chain components reveals a marked deficiency in Tumor 4 (Fig. 6F), suggesting a regulatory defect. Expression of nuclear-encoded oxidative phosphorylation genes is co-regulated and primarily governed by the transcription factors NRF-1 and NRF-2 and the co-activator PGC-1α [74–76]. Both *NRF1* and *NFE2L2*, encoding NRF-1 and NRF-2, respectively, were expressed at low levels throughout the tissue (Supp. Fig. 7). However, *PPARGC1A*, encoding PGC-1α, was absent in Tumor 4 (Fig. 7F, Supp Fig 7). Re-assessment of scDNA-seq reveals LOH at the *PPARGC1A* locus on chr4p specifically in Clone A (Fig 6G). We can therefore hypothesize that a clone-specific *PPARGC1A* inactivating event, such as hypermethylation or an undetected deleterious mutation, promotes an oxidative phosphorylation deficit in those cells.

**Figure 7.**
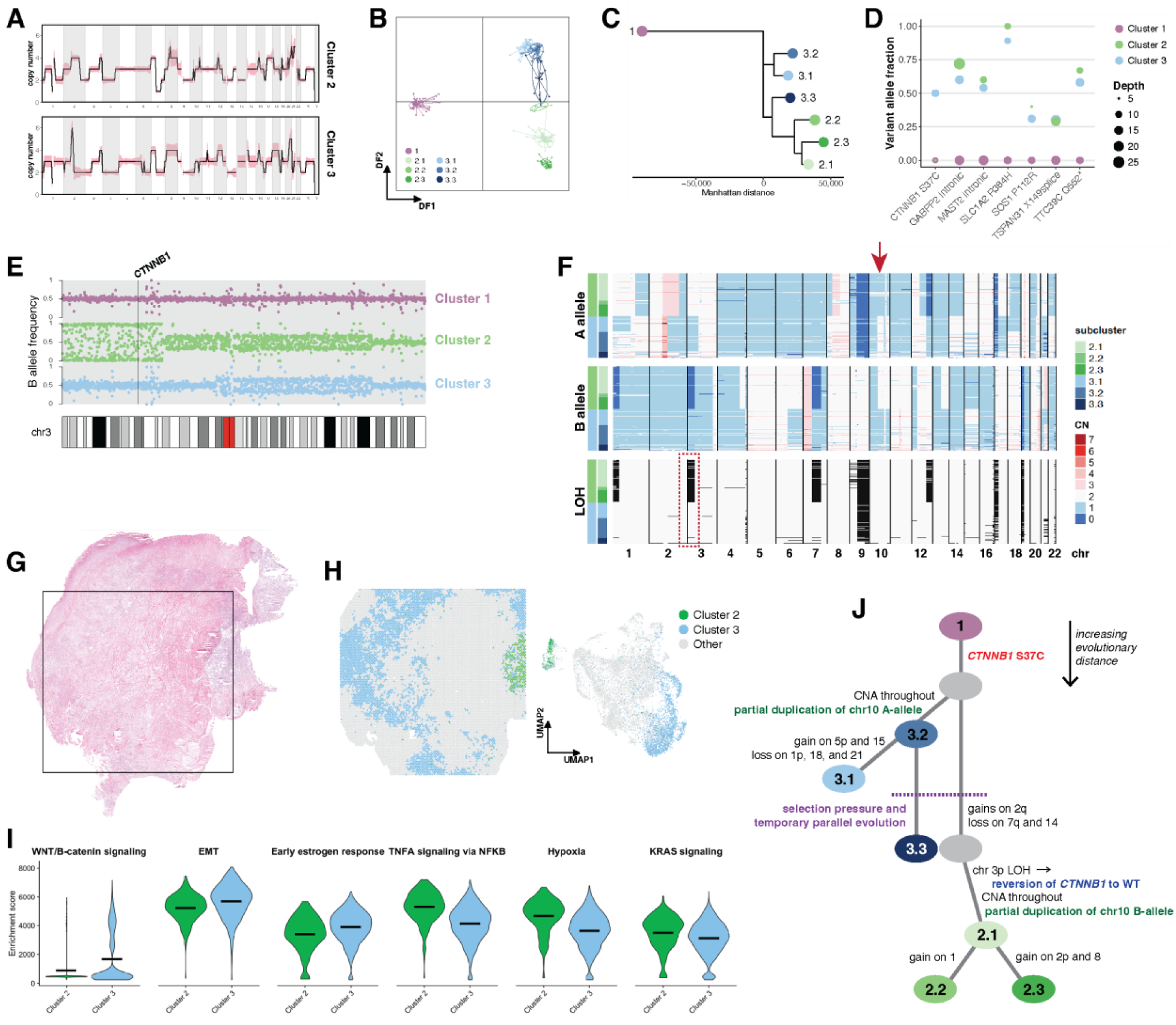
Unique evolutionary history in a multi-clonal CCOC. (A) Average copy number profiles for clusters 2 and 3 of OV511. Shading represents standard deviation. (B) DAPC analysis applied to the subclusters of OV511. (C) Subcluster phylogeny of OV511. Cluster 1 was specified as the outgroup, and is displayed in negative evolutionary space for visualization purposes. (D)Variant allele fractions for select somatic mutations shared by clusters 2 and 3 of OV511. (E) B allele frequency in 100 Kb blocks across chromosome 3 for OV511 clusters 2 and 3. The *CTNNB1* locus is annotated. (F) Single cell allele-specific copy numbers and inferred LOH for sample OV511. LOH is represented by black shading. The arrow marks the location of a mirrored convergent evolution event on chromosome 10. At this region, cluster 2 has two copies of the B allele and one copy of the A allele, while cluster 3 has one copy of the B allele and two copies of the A allele. Both clusters thus have an absolute copy number of 3. The dotted box highlights the LOH on chromosome 3 that results in *CTNNB1* wild-type reversion. (G) H&E stained tissue section of OV511. The square denotes the 11mm x 11mm area captured by the Visium assay. (H) Spatial map and UMAP of ST spots determined to represent clusters 2 and 3 by positional gene set enrichment. (I) Aggregate enrichment scores for ST annotations in (H). The mean of each dataset is annotated. All plots shown have a Bonferroni-corrected p-value < 0.0001 (J) Hypothesized evolutionary trajectory for sample OV511. Somatic *CTNNB1* mutation is the oncogenic driver. Two main branches are distinguished by unique CNA, including mirrored allelic imbalance on chr 10. A parallel evolution event drives similar subclonal alterations, before cluster 2 diverges with a unique set of CNAs, including loss of the *CTNNB1* mutation.

### Unique evolution of a multi-clonal CCOC

Tumor OV511 contained two distinct tumor clones, clusters 2 and 3, with several shared characteristics, including gains of chr5, 7p, and Xq (Fig. 7A, Supp Fig. 8). However, cluster 2 exhibited unique gains of regions of 2q, 4q, and 13q, and loss within 7q. Cluster 3 had gain of 1q, a unique focal amplification within 2q, and gain of chr14. Further heterogeneity is seen within subclusters, with most subclusters having at least two distinguishing variations. Subcluster 3.3 is notable for sharing several characteristics with cluster 2, including gain in similar regions of 2q, loss at 7q, and two copies of chr14 (Fig 2). DAPC confirms the intermediacy of subcluster 3.3, as does phylogenetic inference (Fig. 7B-C).

Variant analysis revealed several shared somatic variants with comparable VAF in both clones (Fig. 7D, Supp. Fig. 9), reflecting a shared ancestry. A *CTNNB1* S37C oncogenic mutation was also identified in cluster 3 (Fig 7D). As cluster 3 is the earlier clone (Fig. 7C), and *CTNNB1* mutations have been identified in CCOC [11, 77], we infer that this is the oncogenic driver mutation. Interestingly, cluster 2 did not harbor the mutation. Assessment of B-allele frequency across chr3 indicates LOH at *CTNNB1* in cluster 2, but not cluster 3 (Fig 7E). Genome-wide allele-specific CN confirms cluster-specific LOH at this location (Fig. 7F), as does manual examination of heterozygous germline SNPs within *CTNNB1* introns (Supp. Fig. 10). This therefore appears to be a spontaneous reversion of an oncogenic mutation in a secondary clone via LOH that retained the wild type allele.

In ST data of OV511, low transcript counts prevented inferCNV from resolving CN profiles. We therefore identified epithelial-majority spots corresponding to clusters 2 and 3 by positional gene set enrichment (Supp. Fig 11). As with other samples, the distinct clones occupied discrete spatial and low-dimensional locations (Fig. 7H) and had distinct histologies (Supp. Fig. 12). Examination of oncogenic pathway expression revealed enrichment of WNT/β-catenin signaling and EMT in cluster 3, consistent with an activating *CTNNB1* mutation (Fig. 7I). This cluster was also enriched in estrogen signaling, which is also linked to β-catenin activity [78]. Cluster 2 had greater enrichment of TNFα and KRAS signaling, as well as hypoxia markers, indicating that these expression programs may compensate for *CTNNB1* reversion in this sample.

We can therefore construct a phylogenetic representation of the unique evolutionary history of OV511 (Fig. 7J). While cluster 3 unambiguously precedes cluster 2, the placement of subcluster 3.3 is confounded by its similarities to cluster 2. From allele-specific CN, we determine that subcluster 3.3 matches cluster 3 in LOH patterns, and in the presence of a mirrored convergent evolution event on chromosome 10 (Fig 7F). Thus, while phylogeny based on absolute copy number alone suggests that subcluster 3.3 and cluster 2 share a most recent common ancestor (Fig. 7C), that is biologically unlikely. Instead, we posit that subcluster 3.3 is evolutionarily grouped with cluster 3, but that the emergence of an environmental pressure selected for similar CNA in subcluster 3.3 and the nascent cluster 2. This temporary parallel evolution diverged when cluster 2 acquired further somatic alterations, including the reversion of *CTNNB1* to wild type.

## Discussion

Ovarian cancer is notable for its high degree of genomic disorder and intra-tumoral heterogeneity. Through integration of spatial transcriptomics and single cell whole genome sequencing, we describe cell-level and subclonal features that would be indistinguishable from each dataset alone.

In two samples, we identify pseudo-diploid cells with non-random, preferential CNA in a diploid majority genome, and confirm the presence of matched somatic mutations in pseudo-diploid and malignant cells. A similar finding was reported by Gao et al in a small number of cells from colon, breast, gastric, and prostate cancers [51]. In the context of ovarian cancer specifically, pseudo-diploid cells may reflect the residual presence of serous tubal intraepithelial carcinomas (STICs), early lesions in the fallopian tube which give rise to HGSOC [79]. STICs have been found to share genomic characteristics with the subsequent tumor, including mutational profiles and amplification of the *CCNE1* locus [80–82]. However, we also report pseudo-diploid cells in a sample of clear cell histology, which is not associated with STIC precursors. Generally, these characteristics of pseudo-diploid cells support the classic model of tumorigenesis, in which somatic alterations are sequential and cumulative [83].

Two multi-clonal tumors provided us the opportunity to infer clonal evolution and associated somatic events. Sample OV594 was a HGSOC with a major clone and a minor clone of significantly fewer cells. After acquisition of an initial, shared *TP53* mutation, the two clones diverged significantly. Each further lost an additional tumor suppressor gene: *SMAD4* in clone A and *PTEN* in clone B. Each clone also had unique regions of gain that were reflected in gene expression differences. In the minor clone, enrichment of transcriptional programs relating to EMT, inflammation, and metabolic capacity correlated with CNAs. Based on both scDNA-seq and spatial transcriptomics, we conclude that the minor clone persisted in the tumor alongside the major clone despite not undergoing major expansion, WGD, or further subclonal development. The restricted expression of *MUC16*, or CA125, to the minor clone also has implications for the effect of tumor heterogeneity on clinical diagnostic testing.

The second multi-clonal sample, OV511, was a CCOC and was comprised of diploid cells and two malignant clones. The early clone harbored a heterozygous *CTNNB1* S37C mutation, a known oncogenic hotspot [84]. Surprisingly, the secondary clone lost this mutation through a loss of heterozygosity event on chromosome 3 that retained the wild-type *CTNNB1* allele. Reversion mutations have been reported previously, but these typically involve restoration of function to tumor suppressor genes [85]. For example, germline *BRCA1/2* mutations may revert when faced with the selective pressure of PARP inhibitor therapies, with which *BRCA* deficiency demonstrates synthetic lethality [86–88]. Recently, loss of an oncogenic driver mutation has been reported in a liver metastasis of an *EGFR*-driven lung adenocarcinoma following treatment with a neopeptide vaccine targeting the mutation [89]. In that case, loss of the *EGFR* mutation was also suspected to be caused by an LOH event.

The reversion of *CTNNB1* in OV511 is unique in that it occurs at the primary site in a treatment-naïve tumor. scDNA-seq data allows us to establish that the two clones of OV511 share a common ancestor based on the following evidence: they share identical copy number gains on several chromosomes, they share identical copy-neutral LOH at several loci – a two-step process unlikely to arise by chance, and they share several hundred somatic passenger mutations. On the premise of a shared ancestral clone, we deduce the order of clone emergence from the mathematical evidence of phylogeny analysis and from the logical evidence of LOH patterns, which can be gained but not lost.

We must therefore speculate on evolutionary reasons for reversion of the *CTNNB1* driver mutation, which provides no obvious fitness advantage. LOH is a common mechanism of tumor suppressor inactivation, and the relevant region on chromosome 3 contains several tumor suppressor genes, including *VHL*, *XPC*, and *MLH1*. While we did not detect any significant mutations in these genes, hypermethylation is common at all three genes in various malignancies [90–92]. Thus, LOH at the region surrounding *CTNNB1* may have resulted in a deleterious effect on tumor suppressor expression. It is also possible that a secondary somatic event elsewhere in the genome allowed the clone to tolerate loss of the initial oncogenic driver; CCOC tumors often have mutations in multiple cancer driver genes [93]. From ST data, we confirm that the secondary clone differs in oncogenic signaling pathways that may compensate for driver mutation loss.

We also note that while reversion of oncogenic driver mutations has rarely been reported, such an event would not be discernable in bulk sequencing. Such reversions may therefore occur more frequently than expected, particularly in tumors prone to CNAs, but remain undetected. Similarly, bulk sequencing would obscure mixed WGD populations and mirrored CNAs. These events highlight the complex evolutionary trajectories that contribute to clonal composition at the time of clinical presentation.

Our study supports the intrinsic role of genomic heterogeneity in the tumor progression model of ovarian cancer, in agreement with other studies that have described the dynamic nature of CNA accumulation within tumors, clones, and subclones [24]. While HGSOC is the EOC subtype most typically associated with chromosomal abberations, we find that CCOC also may harbor complex and consequential genomic events. Examination of single cell genomes is therefore particularly valuable in cancers driven by chromosomal instability, which remain clinically challenging.

## Acknowledgments

Funding for the research project was provided by a generous gift from David Sands; from the Department of Translational Genomics at the University of Southern California; and from the Department of Integrative Translational Sciences at the Beckman Research Institute at City of Hope. Services in support of the research project were provided by the Gynecological Tissue and Fluid Repository at the University of Southern California. Sequencing for the project was partially performed by the Keck Genomics Platform at the University of Southern California, supported, in part, with funding from NIH-NCI Cancer Center Support Grant P30 CA014089.

---

**Supplemental Figure 1.**
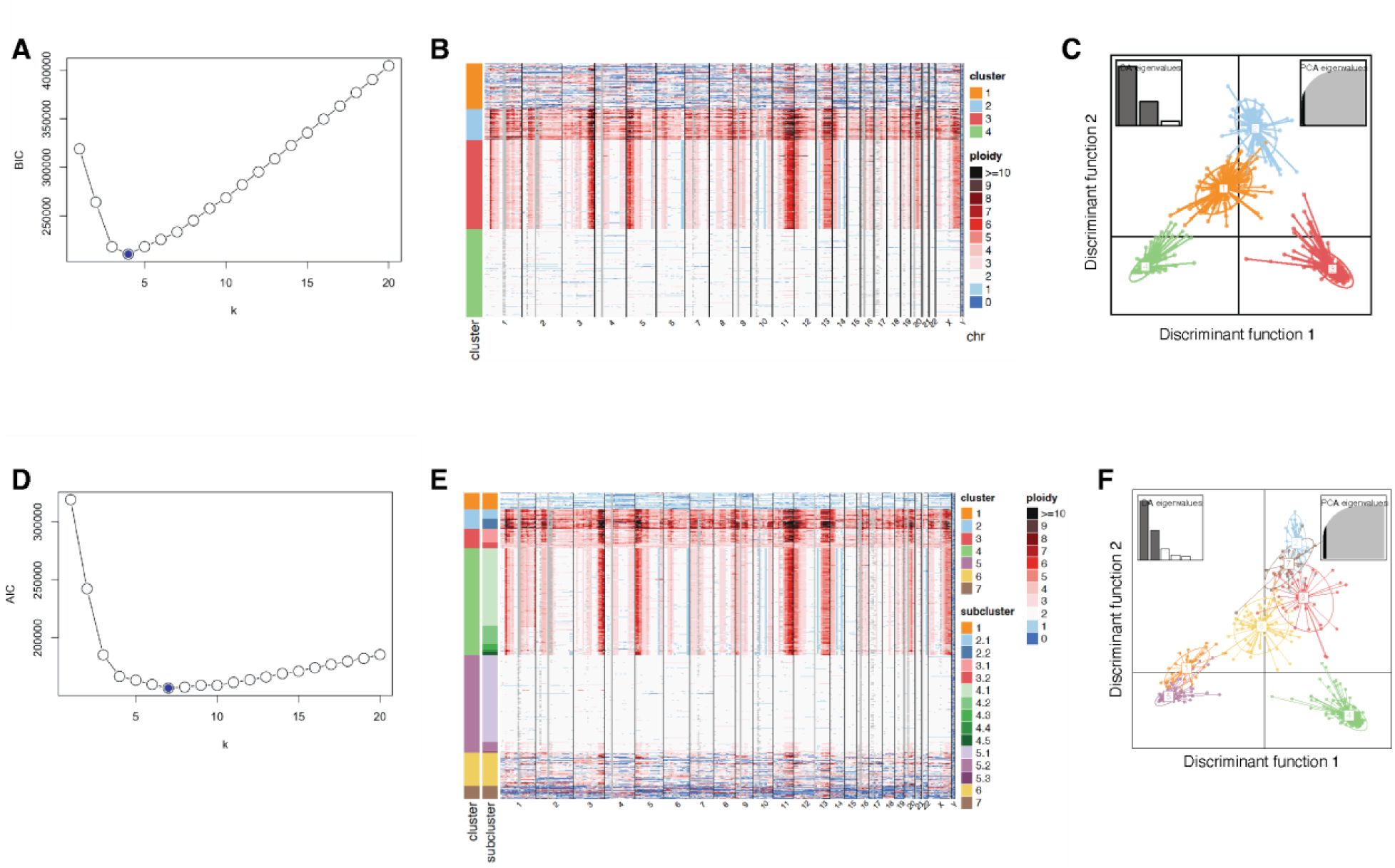
Clustering of scDNA copy number data. (A) Bayesian Information Criterion (BIC) of clustering solutions up to k = 20 for sample OV440. (B) Single cell copy number heatmap reflecting clustering of OV440 based on k = 4, as informed by BIC. (C) Discriminant analysis of principal components applied to k = 4 clusters of OV440. (D) Akaike Information Criterion (AIC) of clustering solutions up to k = 20 for sample OV440. (E) Single cell copy number heatmap reflecting clustering of OV440 based on k = 7, as informed by AIC. Clusters 1, 6, and 7 were excluded from further analysis due to degradation of DNA. (F) Discriminant analysis of principal components applied to k = 7 clusters of OV440.

**Supplemental Figure 2.**
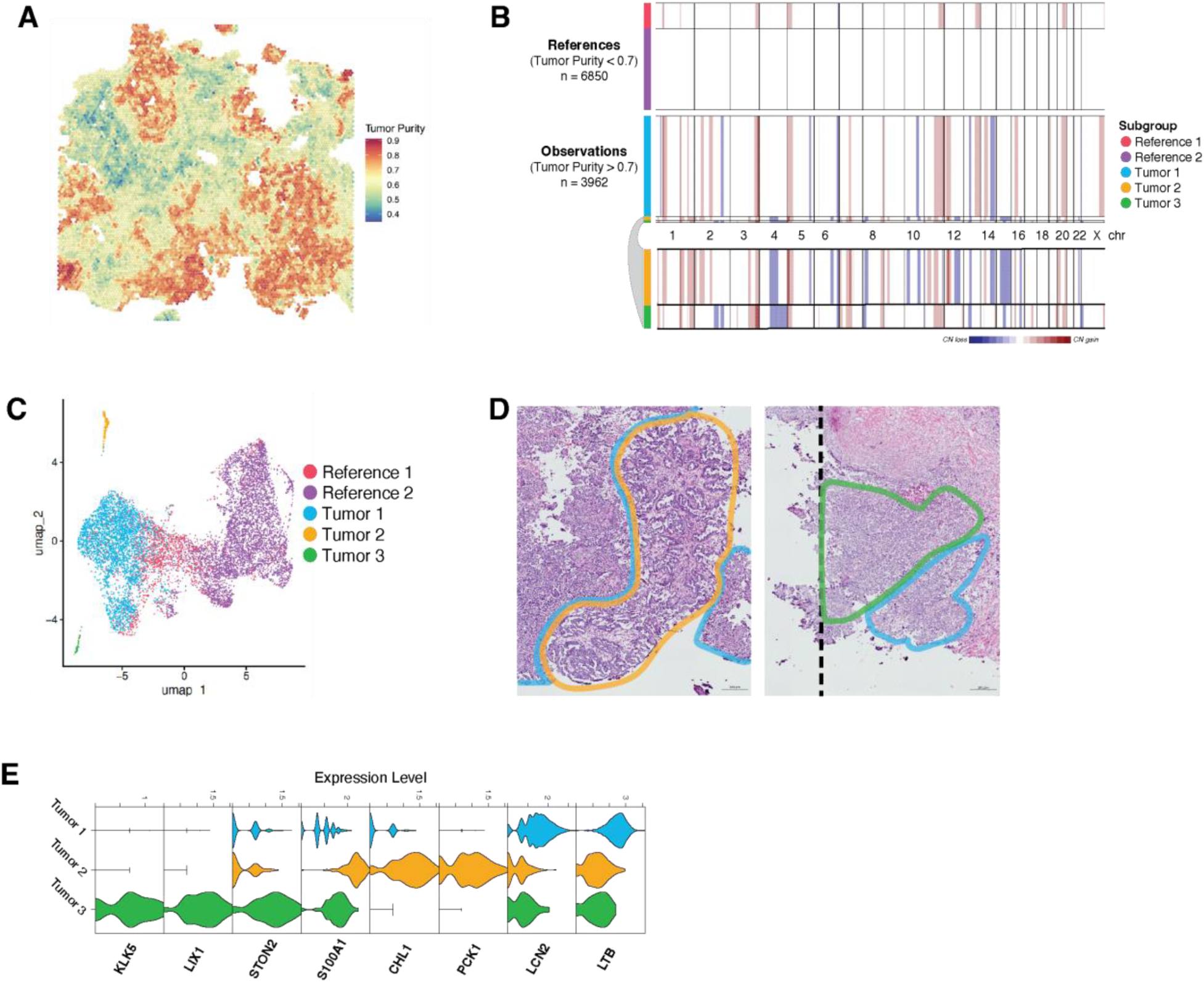
Copy number inferred from OV440 ST data. (A) Tumor purity as determined by ESTIMATE, which was used to determine reference and observation annotations for inferCNV. (B) Full inferCNV results for OV440. Subclusters were determined by Leiden clustering and are annotated on the left bar. HMM-based copy number prediction was performed at the subcluster level. Tumors 2 and 3 are expanded beneath the plot for clarity. (C) Subgroups identified in (C) mapped in low-dimensional space. Reference 2 was determined to likely comprise tumor-diploid mixtures at the tumor-stroma interface. (D) Magnified regions corresponding to the tumor subgroups identified. Tumor 2 displays a papillary growth pattern, tumor 3 displays a micropapillary growth pattern, and tumor 1 displays both micropapillary and solid growth patterns. Scale bar represents 200 microns. (E) Log-transformed gene expression of select genes differentially expressed between clusters.

**Supplemental Figure 3.**
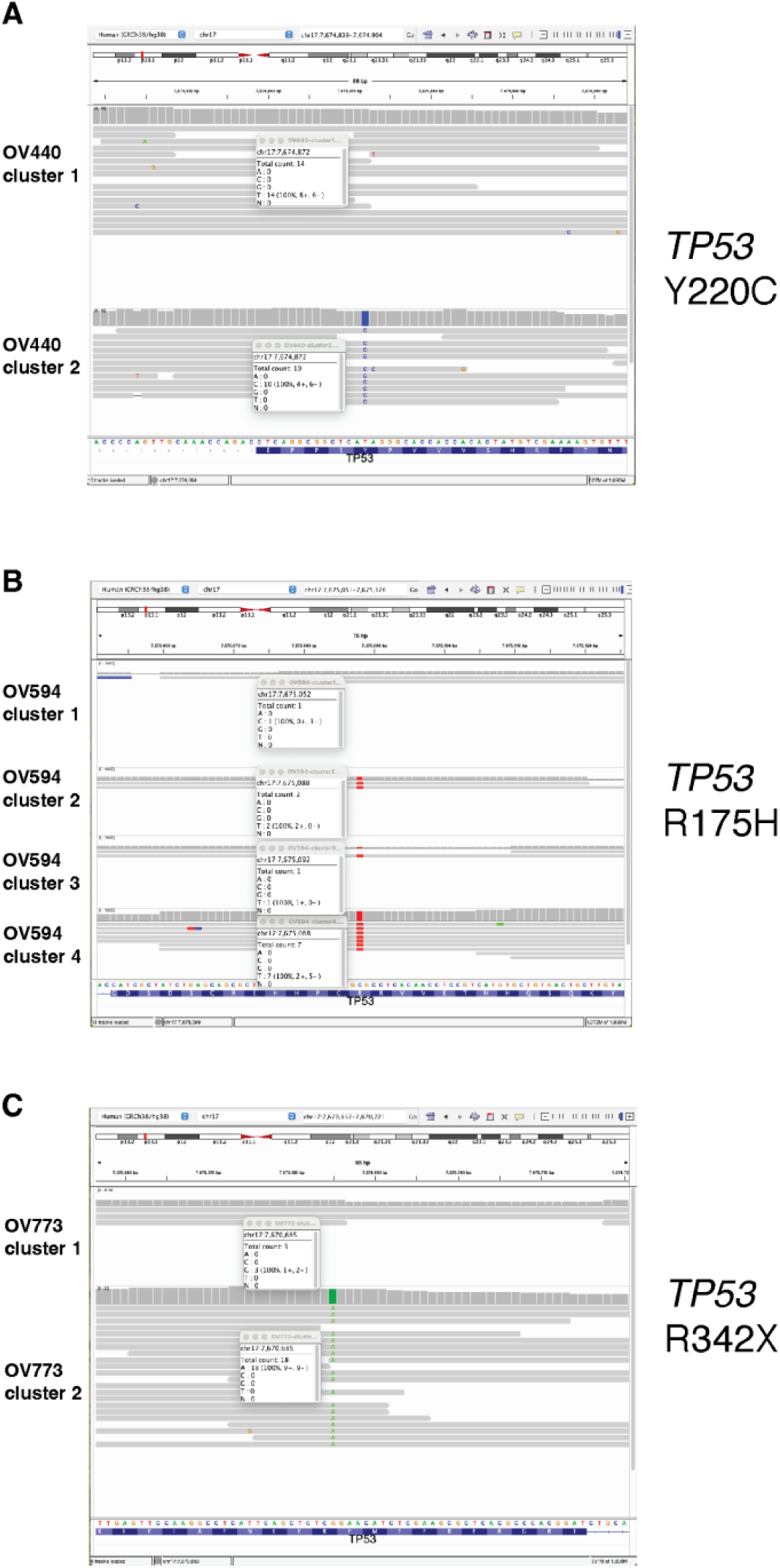
TP53 mutations in HGSOC samples. The clonal *TP53* mutations in samples (A) OV440, (B) OV594, and (C) OV773.

**Supplemental Figure 4.**
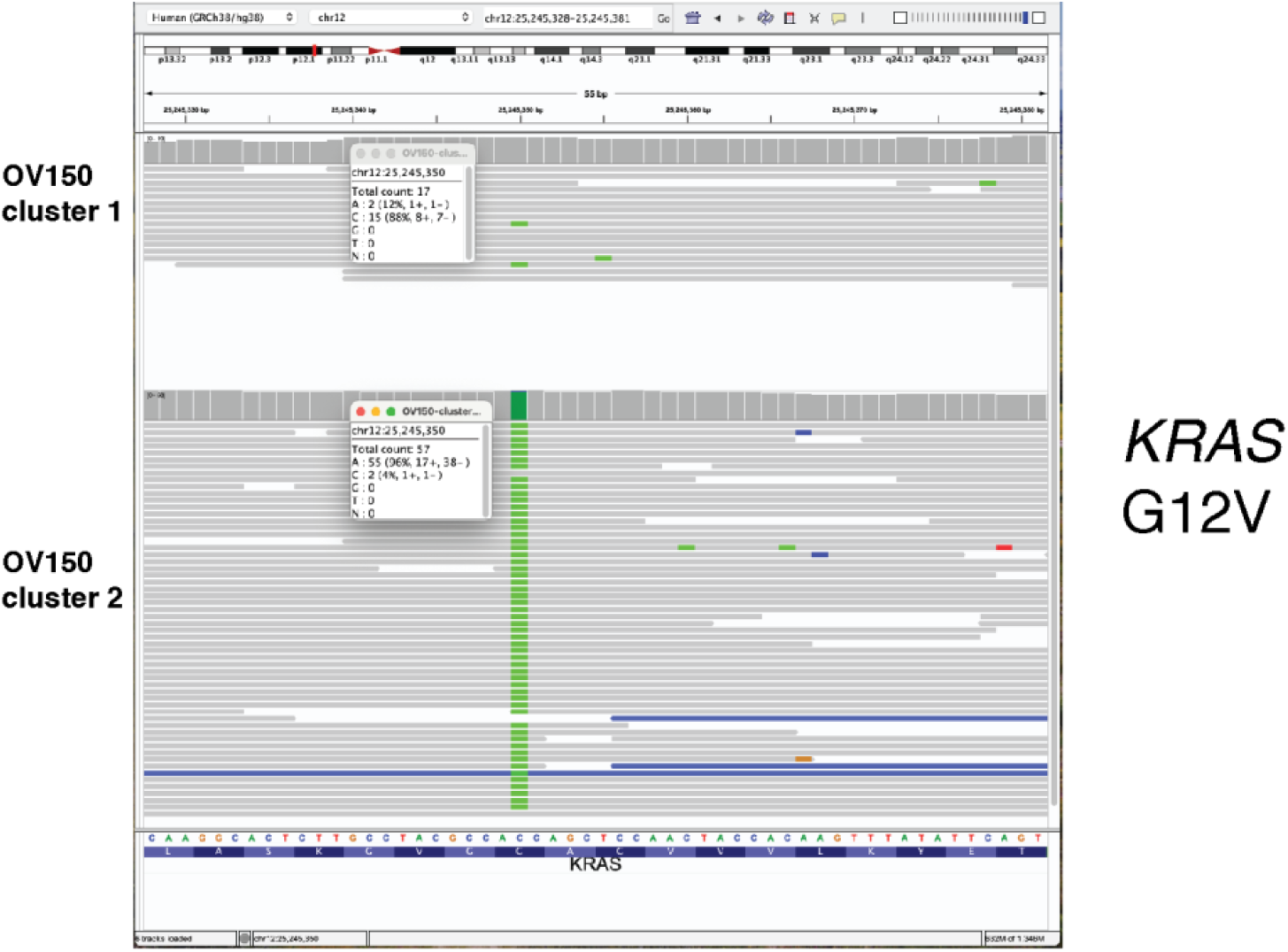
OV150 somatic KRAS mutation. The clonal *KRAS* mutations in OV150 visualized with IGV. The two reads in cluster 1 with the mutation belong to cell 570, which is a pseudodiploid cell described in Figure 4.

**Supplemental Figure 5.**
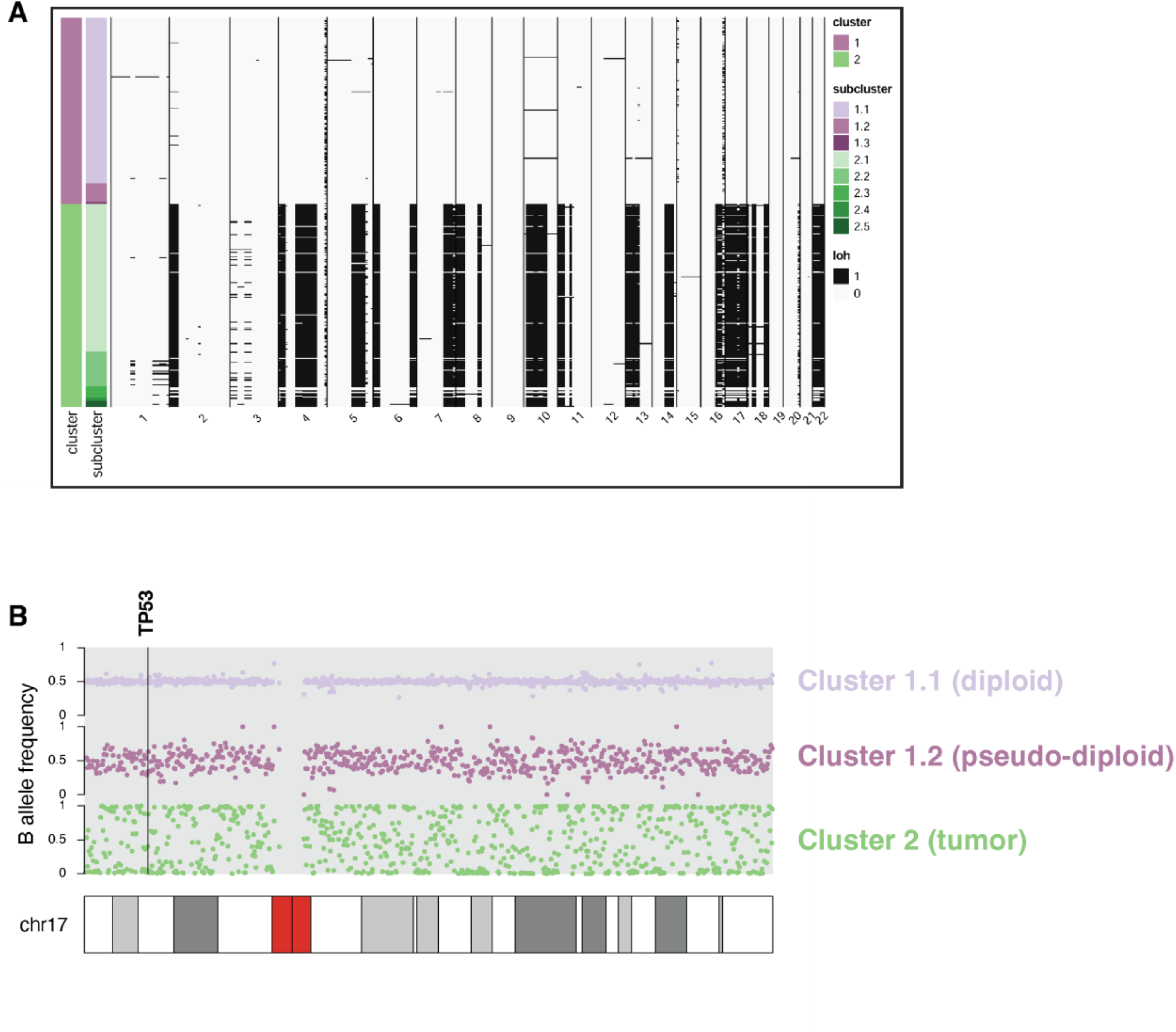
Chromosome 17 LOH in OV440. (A) Loss of heterozygosity in sample OV440 inferred from single-cell allele specific copy number determined by CHISEL. Black shading represents LOH. (B) B allele frequencies across chromosome 17 for clusters of OV440.

**Supplemental Figure 6.**
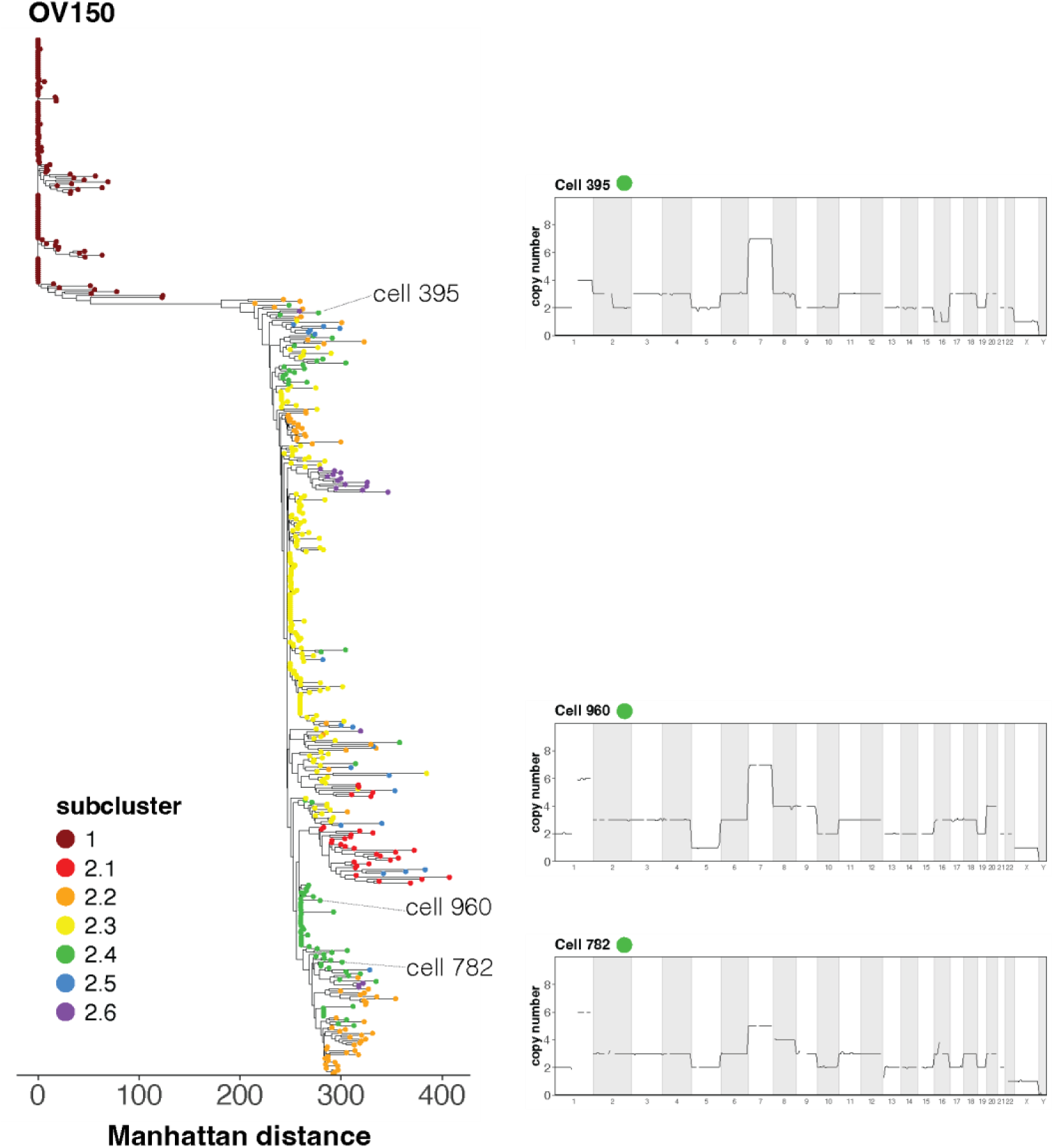
Evolution of OV511 subcluster 2.4. Single cell phylogeny of OV150 (also presented in Figure 4) annotated with three representative cells from subcluster 2.4 and their copy number plots.

**Supplemental Figure 7.**
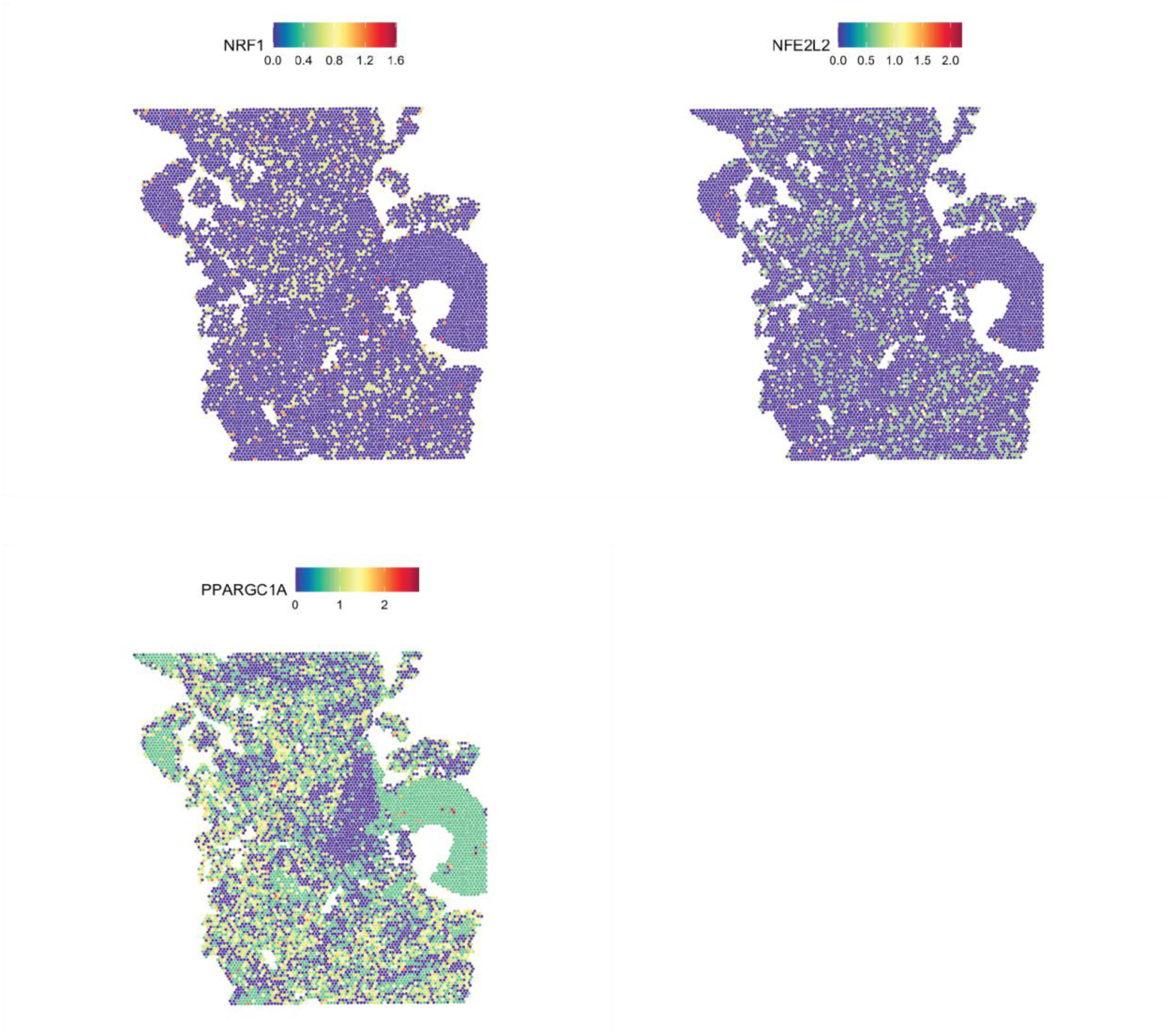
NRF1 and NFE2L2 expression in OV594. Spatially mapped log-transformed gene expression of *NRF1*, *NFE2L2*, and *PPARGC1A* in sample OV594.

**Supplemental Figure 8.**
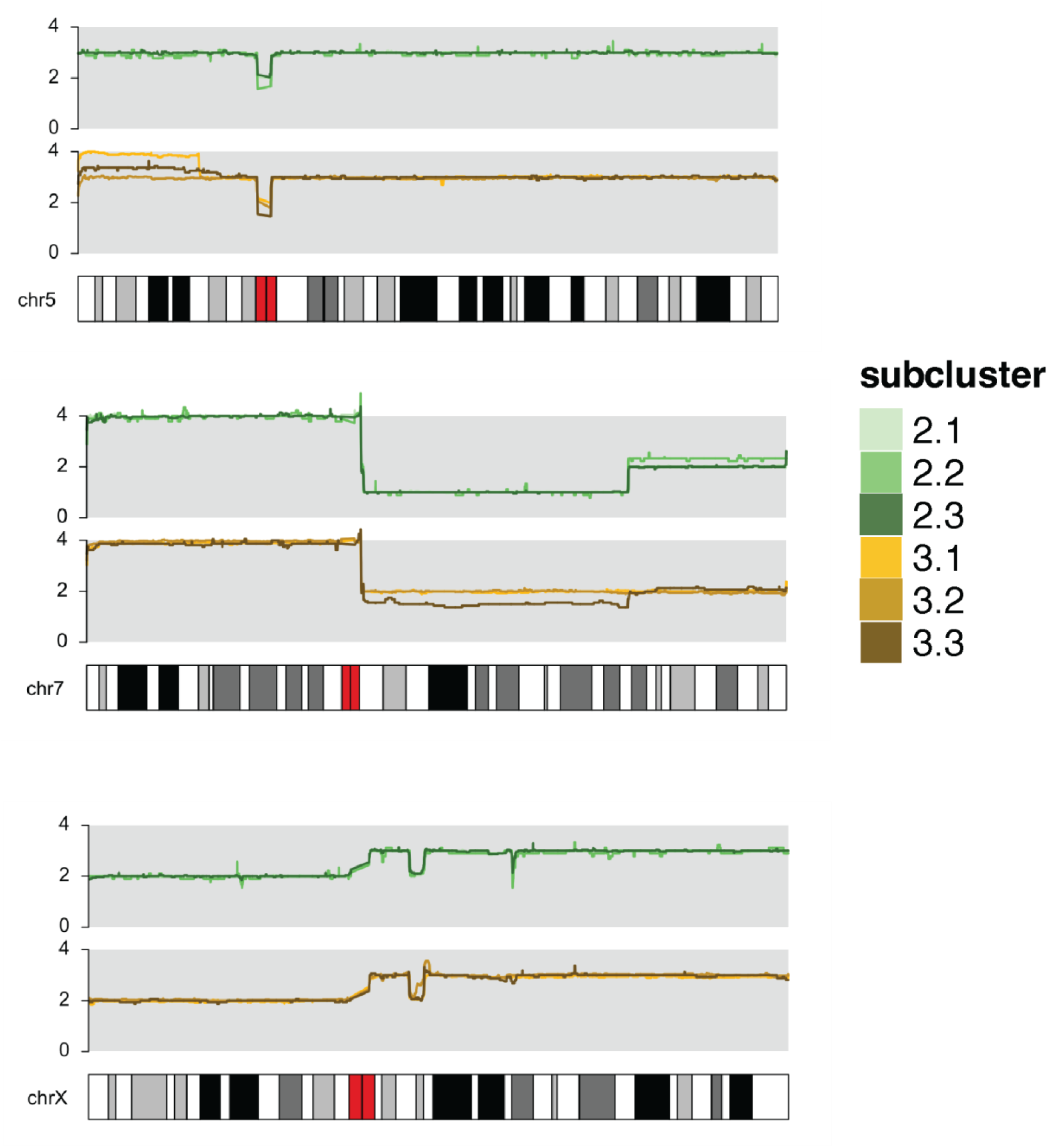
Shared copy number events in OV511 clones. Average copy number line plots for sample OV511, at three representative shared copy number alterations. Copy number is displayed on the y-axis and plotted at 20Kb resolution.

**Supplemental Figure 9.**
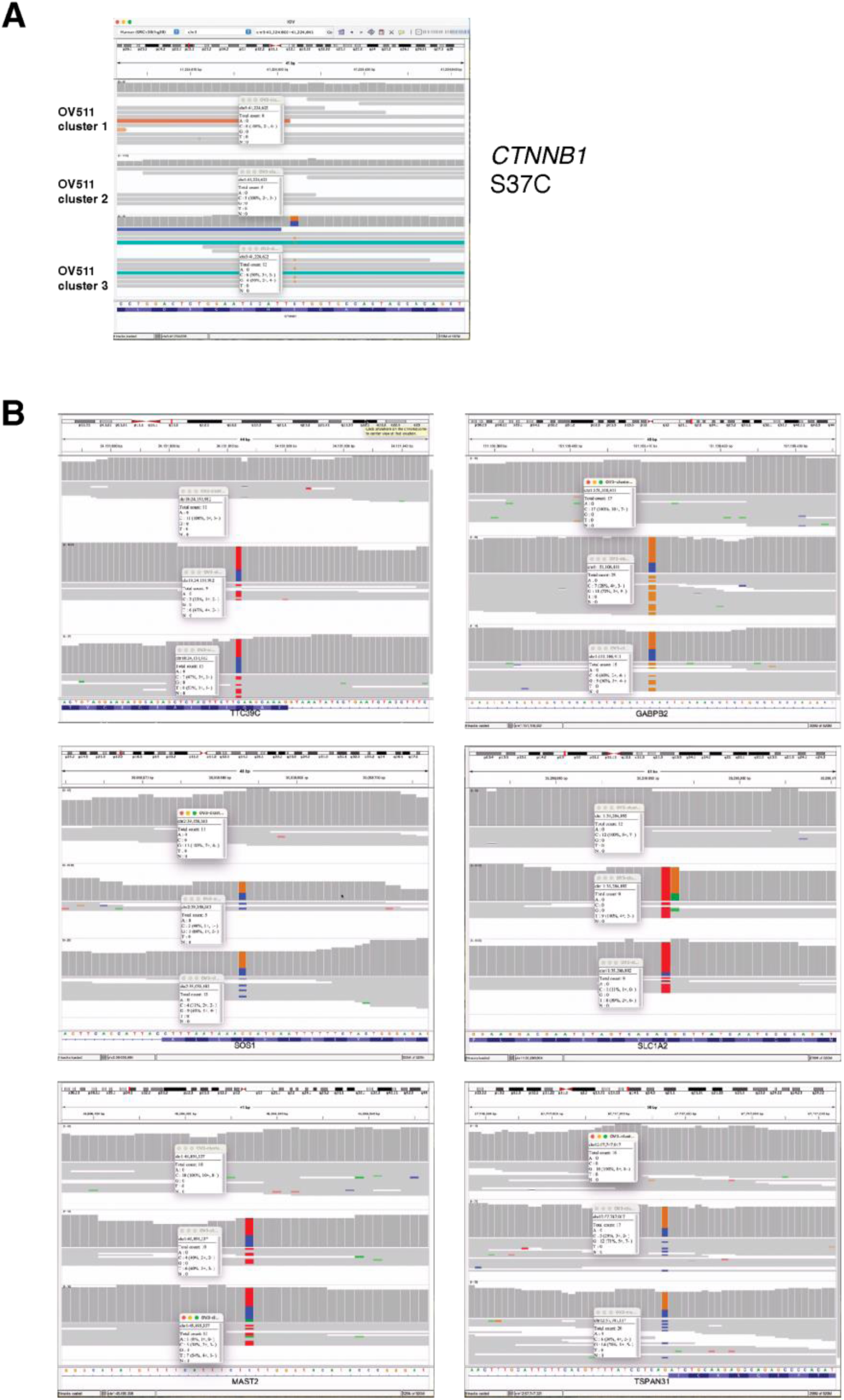
OV511 somatic variants. (A) The putative *CTNNB1* oncogenic driver mutation in sample OV511. (B) Select somatic passenger mutations in sample OV511.

**Supplemental Figure 10.**
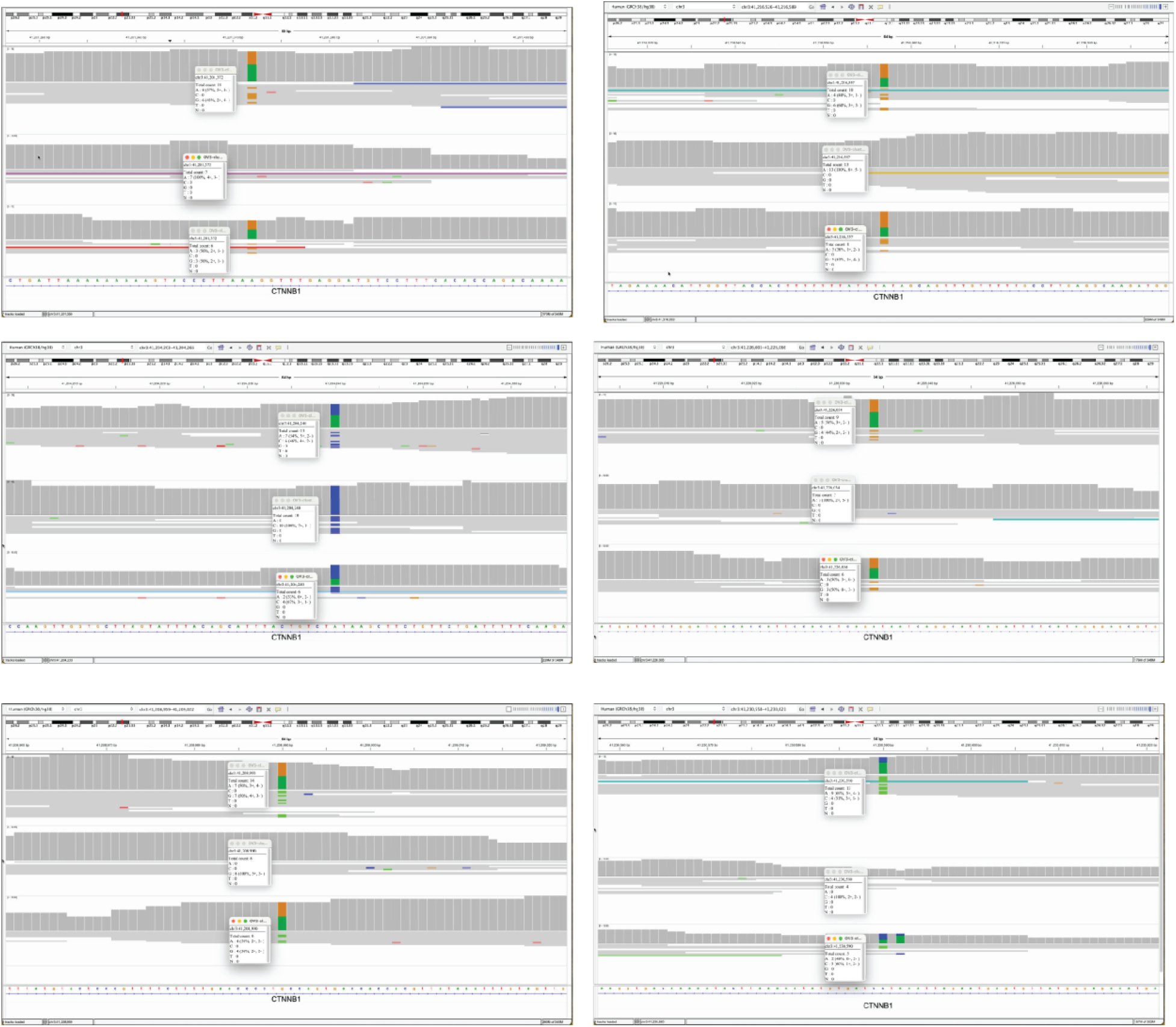
LOH at CTNNB1 SNPs. Heterozygous germline SNPs within *CTNNB1* support LOH in cluster 2 of OV511. Tracks are in order from top to bottom: cluster 1, cluster 2, cluster 3

**Supplemental Figure 11.**
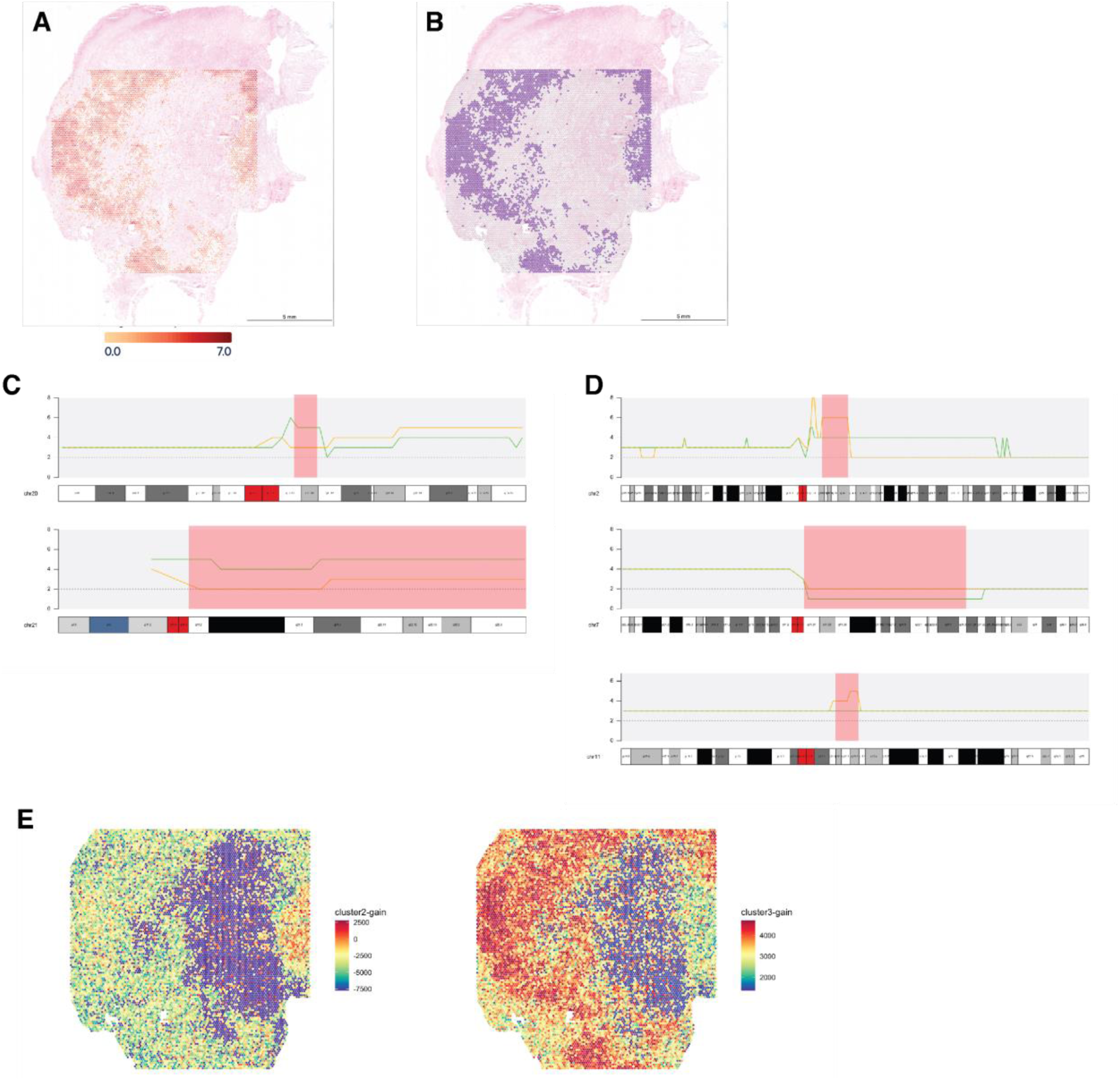
Positional gene set enrichment in OV511. (A) Aggregate expression of *EPCAM*, *KRT7*, and *KRT8* in OV511. Scale bar represents log2 of the summed expression. (B) Spots passing the filtering threshold of log2 summed expression > 1.5. We consider these spots to be comprised mostly of epithelial cells. (C) Regions of the genome utilized in the positional gene set for cluster 2. All unique canonical genes in these regions were exported from UCSC genome browser. (D) Regions of the genome utilized in the positional gene set for cluster 3. (E) Spatially mapped ssGSEA enrichment scores corresponding to the custom positional gene sets for each cluster. Spots passing epithelial filtering (B) were assigned to cluster 2 if the cluster 2-gain score was greater than 2500, and to cluster 3 if the cluster3-gain score was greater than 3000.

**Supplemental figure 12.**
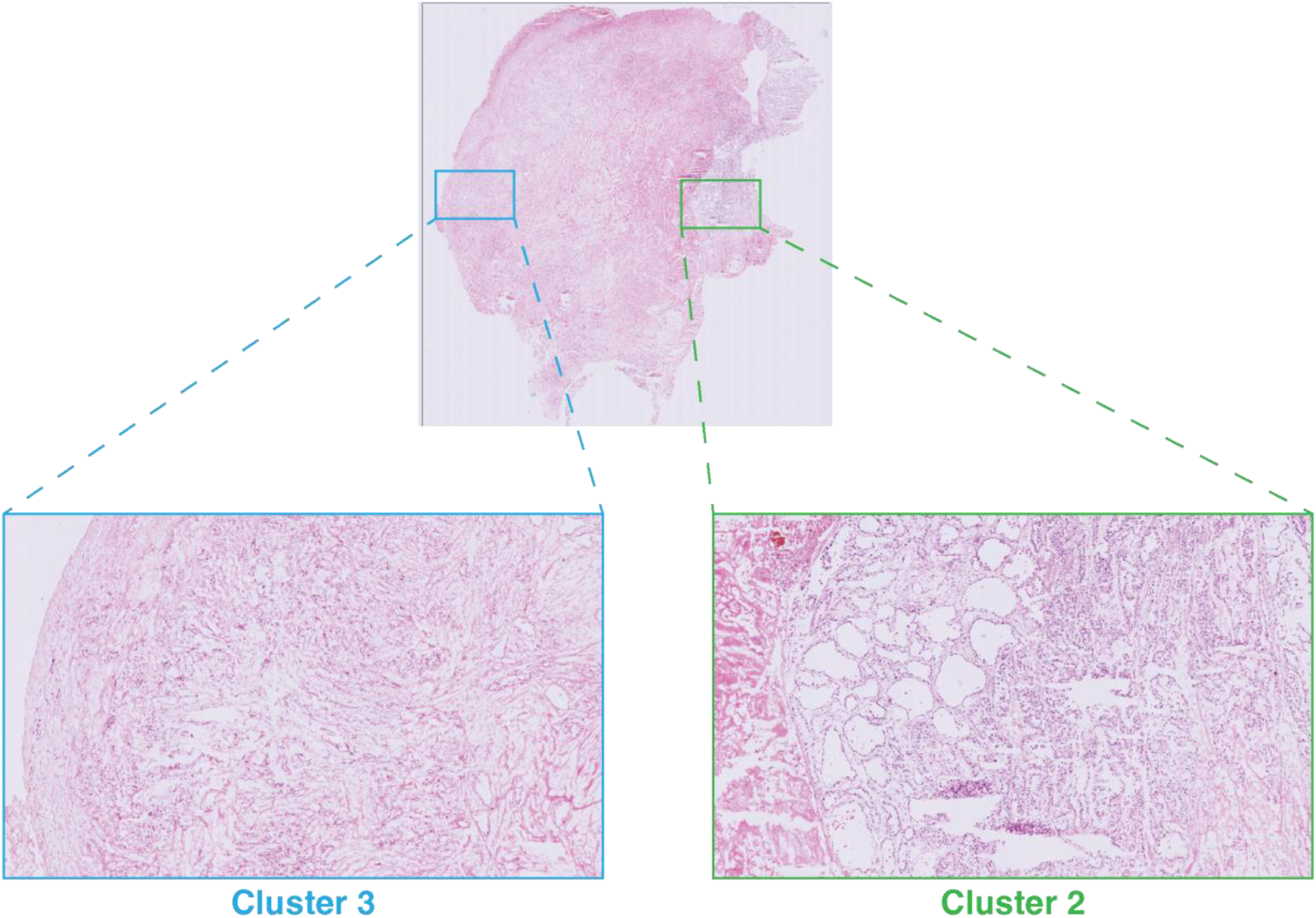
Histology of OV511 clones. Magnified H&E-stained tissue from sample OV511. The indicated regions were determined to correspond to the labeled clusters by positional gene set enrichment.

